# Distinct trajectories of urbanization shape the human gut microbiome across South Asia

**DOI:** 10.64898/2026.01.22.701183

**Authors:** Shreya L. Ramachandran, Nagarjuna Pasupuleti, Richard J. Abdill, Gamini Adikari, Bhavna Ahlawat, Ran Blekhman, Victoria Burgo, Emily R. Davenport, Aparna Dwivedi, Shristee Gupta, Thadiyan Parambil Ijinu, Tiatoshi Jamir, Roheet Karu, Amit Kaushik, John Milton Komarabathini, Laithangpuii, Joseph Lalzarliana, Tsering Norbu, Tlana Pautu, Kantikumar Pawar, Arjun S. Raman, Ramanujam Ramaswamy, Ruwandi Ranasinghe, Rashmi, Sreejith Pongillyathundi Sasidharan, Tharigopula Satheesh, Shalini Shaji, Jason W. Shapiro, Ekta Singh, Vanlalhuma Singson, Anitha Sundararajan, Kamani H. Tennekoon, David Tetso, Jose A. Urban Aragon, Anjana H.J. Welikala, Ruokuonuo Rose Yhome, Niraj Rai, Maanasa Raghavan

**Affiliations:** Department of Human Genetics, University of Chicago, Chicago, IL 60637, USA; Birbal Sahni Institute of Palaeosciences, Lucknow, Uttar Pradesh 226007, India; Section of Genetic Medicine, Department of Medicine, University of Chicago, Chicago, IL 60637, USA; The Postgraduate Institute of Archaeology (PGIAR), University of Kelaniya, Colombo 00700, Sri Lanka; Indian Institute of Technology (IIT) Gandhinagar, Gandhinagar, Gujarat 382355, India; Duchossois Family Institute, University of Chicago, Chicago, IL 60637, USA; Department of Biology, Huck Institutes of the Life Sciences, The Pennsylvania State University, University Park, PA 16802, USA; Naturae Scientific, Kerala University-Business Innovation and Incubation Centre, Kariavattom Campus, Thiruvananthapuram 695581, Kerala, India; The National Society of Ethnopharmacology, VRA 179, Mannamoola, Peroorkada P.O, Thiruvananthapuram 695005, Kerala, India; Academy of Bharatiya Knowledge Systems and Traditions, VRA 179, Mannamoola, Peroorkada P.O, Thiruvananthapuram 695005, Kerala, India; Nutraceuticals-India Consortium, Naturae Science Foundation, 21 NCC Nagar, Peroorkada P.O, Thiruvananthapuram 695005, Kerala, India; Department of History & Archaeology, Nagaland University, Kohima Campus, Meriema, Nagaland 797004, India; VARASA-Association of Cultural Heritage and Archaeology, Pune, Maharashtra 411038, India; Chaudhary Charan Singh Haryana Agricultural University, Hisar, Haryana 125004, India; Government Degree College, Bhadrachalam, Andhra Pradesh 507111, India; Department of History, ICFAI University, Aizawl, Mizoram 796015, India; Department of History and Ethnography, Mizoram University, Aizawl, Mizoram 796004, India; Rock Art and Historical Society of Spiti, Kaza, Himachal Pradesh 172114, India; Independent researcher affiliated with SEL Foundation, Aizawl, Mizoram, India; Archaeological Survey of India, Aizawl, Mizoram 796001, India; Department of Pathology, University of Chicago, Chicago, IL 60637, USA; Center for the Physics of Evolving Systems, University of Chicago, Chicago, IL 60637, USA; Institute of Biochemistry, Molecular Biology and Biotechnology, University of Colombo, Colombo 00300, Sri Lanka; Multidisciplinary Research Unit (Department of Health Research, MoHFW, GoI),Government Medical College, Thiruvananthapuram 695011, Kerala, India; Department of Research, Vivekananda Memorial Hospital, Swami Vivekananda Youth Movement, Hanchipura Road, Saragur, Saragur Taluk, Mysuru, Karnataka 571121, India; Department of Anthropology, University of Hyderabad, Gachibowli, Hyderabad, Telangana 500046, India; Center for Research Informatics, University of Chicago, Chicago, IL 60637, USA; Department of Earth Sciences, Indian Institute of Technology, Roorkee, Uttarakhand 247667, India; Art and Culture Department, Government of Mizoram, Aizawl, Mizoram 796007, India; Department of Anthropology, Kohima Science College, Kohima, Nagaland 797002, India; The Highland Institute, Kohima, Nagaland 797003, India; Committee on Genetics, Genomics and Systems Biology, University of Chicago, Chicago, IL 60637, USA; Committee on Southern Asian Studies, University of Chicago, Chicago, IL 60637, USA

**Author notes:** Co-corresponding authors. Maanasa Raghavan; Niraj Rai. These authors contributed equally (Shreya L. Ramachandran, Nagarjuna Pasupuleti). These authors are listed alphabetically.

## Abstract

Human gut microbiomes respond to lifestyle transitions, yet the extent to which these responses are conserved across spatio-cultural contexts remains undercharacterized. We present the South Asian MicroBiome ARray (SAMBAR), a population-scale 16S gut microbiome study of 575 adults from ten geographically and socio-culturally diverse South Asian communities. Each community was sampled in ancestral villages and urban centers, enabling controlled comparisons of geography and lifestyle. Relative to global cohorts, SAMBAR microbiomes occupy a distinct compositional space with stronger correlation to geography and community membership than lifestyle. Although urbanization is consistently associated with increased abundance of disease-linked taxa, microbiome responses to lifestyle transitions are largely community-driven, including the acquisition of wheat– and dairying-associated microbial modules in some communities that may facilitate non-genetic adaptation to lactase non-persistence. Microbiome responses to urbanization are heterogeneous even at regional scales, reflecting local culture and geography and underscoring the need for community-specific investigations of health impacts.

## Introduction

Around 10-12 thousand years ago, human groups in the Fertile Crescent began a transition from a hunting and gathering subsistence strategy to a lifestyle based on agriculture, a process that occurred independently and at varying times in other regions of the world^1^. Since then, the transition to agro-pastoralism has, in large part, swept the world; yet, there remain some populations today that continue to adhere to a hunter-gatherer subsistence strategy, primarily found in regions of Africa, South Asia, and South America. However, for many, their traditional way of life is rapidly being replaced by industrialization and the adoption of agricultural and/or processed food techniques^2,3^. This is in part due to rapid globalization and increased immigration in the last century, leading to the spread of a Westernized or urbanized diet that is typically characterized by higher levels of fat, simple sugars, and processed foods and is also linked with compositional microbial changes, decreased diversity of the microbiota, and health outcomes^4–6^.

Previous studies have found differences between the microbiomes of hunter-gatherer and urbanized populations. Taxa associated with these lifestyles are often described as VANISH (volatile and/or associated negatively with industrialized societies of humans) and BloSSUM (bloom or selected in societies of urbanization-modernization)^7^. For example, the microbiomes of the Hadza hunter-gatherers in Tanzania showed an overall increased microbial diversity as well as significant differences in the abundances of specific taxa compared to European control populations^8^. More recent work across four societies at different points of lifestyle transition in Himalayan populations identified differences across populations and correlations between lifestyle and taxonomic profiles, but unlike the Hadza studies, decreases in microbiome diversity were not observed as populations became more Westernized ^9^. These studies strongly suggest that transitioning to agro-pastoralism has significant effects on the gut microbiome, but the precise mechanisms may be population-specific and necessitate more extensive and diverse microbiome surveys at regional scales. Diversifying the microbiome literature is critical given European and North American donors have continued to represent over 50% of the global datasets in public repositories such as the Sequence Read Archive over the past decade^10^.

Several South Asian Indigenous/tribal populations are rapidly transitioning away from their traditional lifestyles due to forced displacement and socio-economic mobility^11–13^. India alone is home to more than 4,500 anthropologically defined populations—including 500 tribes, of which 72 are traditionalist hunter-gatherers^14^ who have already transitioned to an urbanized lifestyle or are currently undergoing transition. Sri Lanka is home to the Indigenous Adivasi population who today live in fragmented settlements across the island and are also undergoing urbanization^15^. Despite the staggering demographic and cultural diversity, there are limited regional microbiome surveys of South Asians, in general, and particularly of transitioning, traditional populations. South Asia’s underrepresentation in global microbiome research is underscored by the Human Microbiome Compendium (v.1.1.1)^16^. Despite South Asians making up approximately 25% of the global population^17^, only 3.3% of the Compendium samples (5,529/168,464) originate from South Asia – an expansive region represented solely by India and Bangladesh. Notably, almost all the samples in the Compendium from South Asia are from studies on infants and children^16,18,19^. Furthermore, numerous public health studies have identified a rising prevalence of metabolic disease in primarily urban South Asians^20–23^. Given the differences in the microbiome profiles of diabetes patients and healthy controls^23,24^, it is critical to investigate the role of the microbiome in shaping the health of South Asians undergoing lifestyle transition. Previous studies in the area have shed critical light on regional microbiomes, but their focus on smaller or geographically mismatched urban and rural cohorts, often with limited lifestyle and phenotype metadata^25–32^, makes their data less relevant to analyses of the ongoing urbanization in the region. Through the implementation of a sampling design grounded in anthropological genetics – consisting of community-matched rural and urban cohorts to ensure human genetic and cultural concordance – this study provides critical insights into the biological and health impacts of lifestyle transition on the microbiomes of vulnerable populations and, more broadly, into the modern-day health landscape of South Asians. We call this dataset the South Asian MicroBiome ARray, or SAMBAR, which is also a lentil and vegetable stew popular in India and Sri Lanka.

## Results

### Indian and Sri Lankan 16S gut microbiome data expands the number of currently available South Asian microbiomes

After our recruitment efforts and informed consent process outlined in Methods, we enrolled individuals from 10 communities that represent a sub-sampling of the linguistic, geographic, and genetic diversity present in India and Sri Lanka (see Methods). In North India, we recruited high-altitude residents of Spiti Valley (>4,000 meters above sea level) in Himachal Pradesh (hereafter, Spitian); in Central India, we enrolled members of the Gond, Kolam, and Koya; in South India, the Kani; and in Northeast India, the Mizo and Pochury Naga (hereafter, Pochury). Within Sri Lanka, we recruited members of the Adivasi community, as well as urban Sinhalese and Sri Lankan Tamil individuals^33^ (Figure 1). These communities, with the exception of the Sinhalese and Sri Lankan Tamil, still practice Indigenous/tribal lifestyles that are characterized by either small-scale foraging and hunting or, in the case of the Spitian individuals, traditional yak herding. However, with improvements in infrastructure and travel connections facilitating access to their ancestral homelands, urbanization has started to impact many of the rural community members at varying rates.

**Figure 1:**
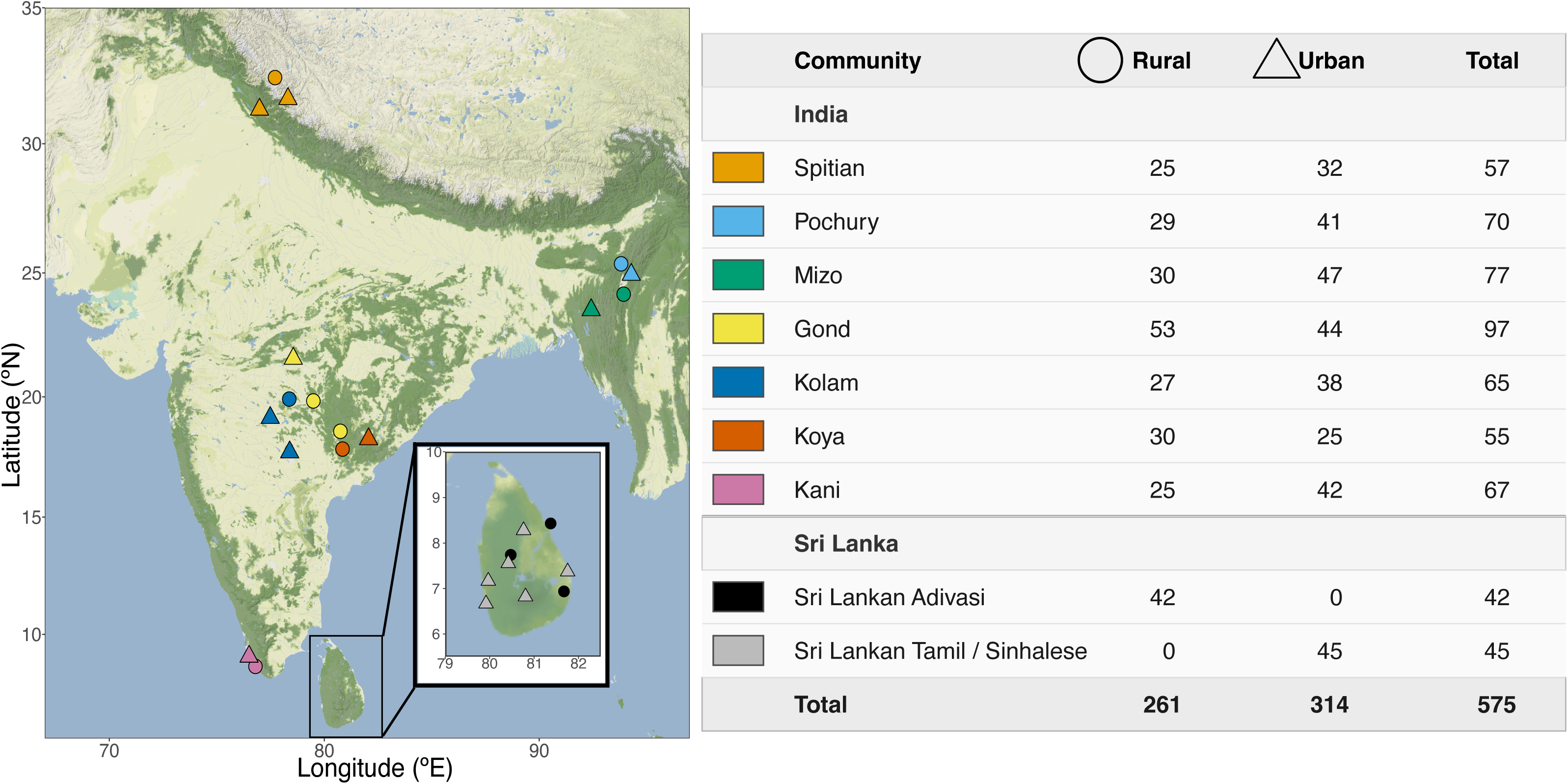
Geographic locations of sampling sites in India and Sri Lanka. Each point on the map represents one district (circle: rural; triangle: urban). Table shows number of urban and rural individuals from each community.

For each of the communities within India, we enrolled participants living in rural settings and practicing more traditional ways of life (‘rural’ cohort) as well as members of the same community who had spent more than a year living in a nearby urban center to their ancestral village (‘urban’ cohort) to ensure matched host genetic backgrounds. In Sri Lanka, because the Adivasi do not have a substantial presence in urban settings, we enrolled the genetically-related Sinhalese and Sri Lankan Tamil^33^ as their urban match. The breakdown of community membership and urban/rural status for the SAMBAR dataset are seen in Fig. 1, with the full description in Tables S1 and S2. In addition to the microbiome sampling, we conducted a detailed diet and lifestyle survey for each participant, inspired by Jha et al^9^. The survey consisted of 36 dietary questions, such as drinking water source, dairy and wheat consumption frequency, and fermented food. Additionally, we recorded demographic data such as language, residence, and occupation, and collected metabolism-relevant phenotypes such as blood pressure, blood glucose, and body mass index (Table S1). This forms an important data resource to capture the variation in lifestyle as it relates to gut microbiome composition, especially because the urban-rural dichotomy likely obscures meaningful differences in each community’s position along the urbanization gradient. At the end of the recruitment process, we enrolled 609 individuals. After quality control and filtering of both the microbiome data and survey data, the final dataset for analysis consisted of 575 individuals. As such, the SAMBAR microbiome and survey dataset makes publicly available more than 500 samples from adults across India, and, for the first time, Sri Lankan individuals spanning the range of communities and lifestyles within the country.

### Differentially abundant taxa in SAMBAR compared to global diversity

To situate the SAMBAR microbiomes within the context of global diversity, we built four size-matched regional cohorts by subsampling from the Human Microbiome Compendium. We selected samples only from studies on adults that used fecal samples and used an amplicon overlapping the V4 hypervariable region. After these filters, four world regions retained enough individuals to randomly sample 575 individuals: Eastern and South-Eastern Asia, Europe and Northern America, Northern Africa and Western Asia, and Sub-Saharan Africa (Data File S1). While South Asia is represented in the Compendium, the vast majority of individuals in the dataset are infants and children with only 30 adult individuals from the region. The SAMBAR dataset increases almost twenty-fold the number of adult South Asian individuals in the dataset.

Compositional microbiome data for all cohorts was centered-log-ratio transformed, and Principal Components Analysis (PCA) was performed on the transformed data (i.e. Aitchison PCA). While the first principal component (PC) largely reflected the Shannon diversity of the dataset (Fig. S1A-B, Table S3), the second and third PCs showed a clear separation of samples by geography (Fig. 2A-B). The second PC broadly separates the SAMBAR and Sub-Saharan African cohorts from the other three regional cohorts, and the third separates SAMBAR from Sub-Saharan Africa. This clustering was statistically significant (PERMANOVA p = 0.001, Table S4-5.) Taxa contributing to PC2 and more prevalent in SAMBAR and Sub-Saharan Africa include taxa canonically associated with rural lifestyles, including Prevotellaceae^34,35^, *Succinivibrio*^36,37^, and *Leyella*^25,38–40^. Taxa contributing specifically to the region of the PCA where SAMBAR samples cluster (Fig. 2A-B) include *Holdemanella, Catenibacterium,* and *Ligilactobacillus* (Fig. S1C).

**Figure 2:**
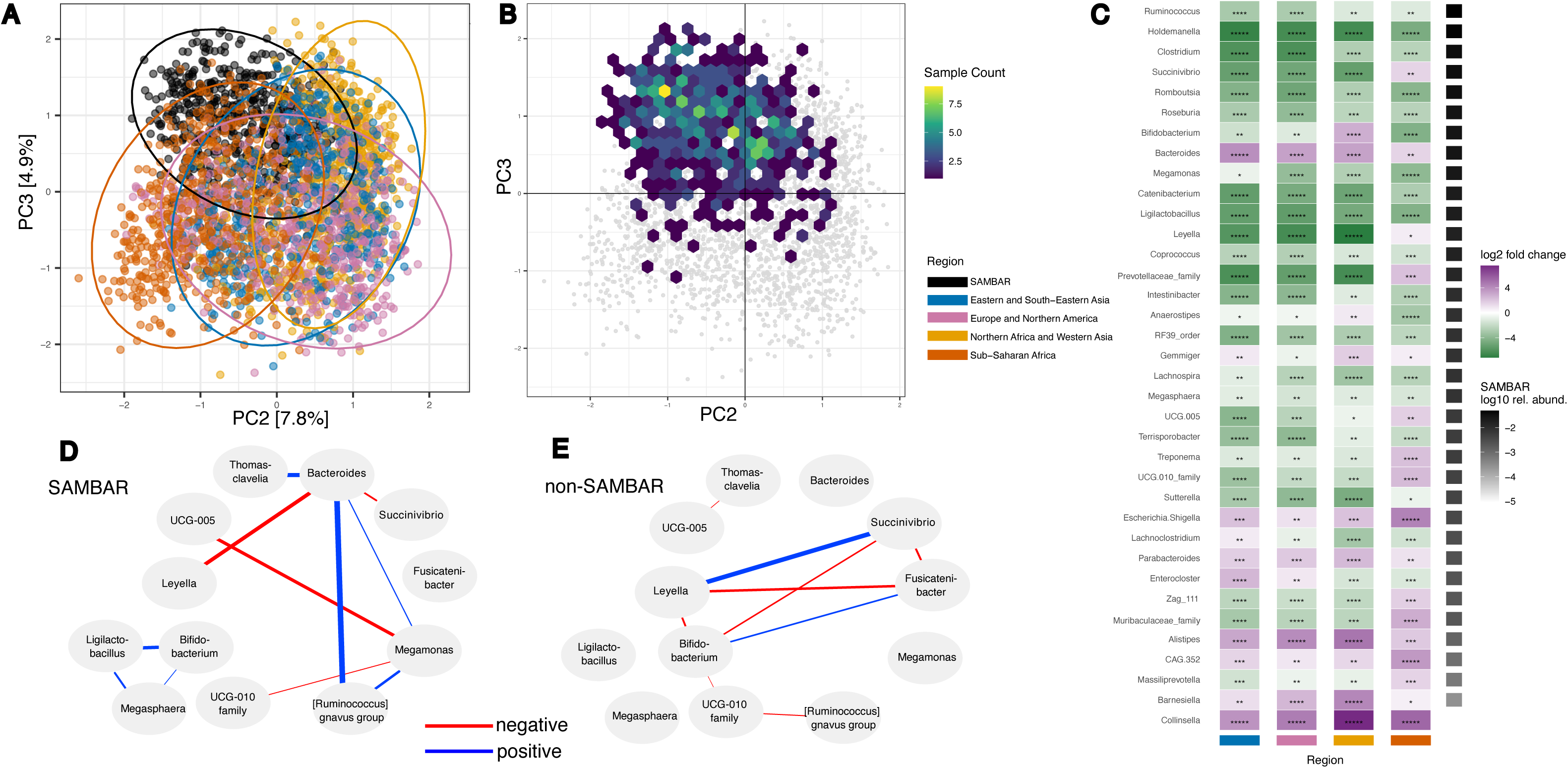
A: Principal Components Analysis (Aitchison distance) based on microbiome composition of SAMBAR cohort combined with four world region cohorts from Human Microbiome Compendium. B: Heatmap showing plotted locations of SAMBAR samples within the PCA shown in 2A. Yellow areas represent a higher concentration of SAMBAR individuals. C: Heatmap showing differential abundance of top 40 most significant taxa between SAMBAR and each of four global cohorts, sorted by descending prevalence within SAMBAR. Green taxa are more abundant in SAMBAR; purple taxa are more abundant in the comparison cohort than in SAMBAR. Taxa are ordered by relative abundance within SAMBAR, represented by black bars. Extended results are shown in Fig. S3. D: Subset of correlation network showing correlations only seen in the SAMBAR dataset. Blue connections signify positively correlated taxa, while red connections reflect negative correlations. Extended results are shown in Fig. S4. D: Subset of correlation network showing correlations only seen in the non-SAMBAR dataset. Blue connections signify positively correlated taxa, while red connections reflect negative correlations. Extended results are shown in Fig. S5.

The PCA biplot visually suggests that specific genera may be enriched or depleted in SAMBAR compared to other world regions. To quantitatively identify these, we used MaAsLin2 to model the differential abundance of taxa between SAMBAR and each size-matched regional cohort drawn from the compendium (Fig. 2C; Table S6; Fig. S3). Taxa significantly enriched in SAMBAR compared to all four regional cohorts are *Ruminococcus, Clostridium, Romboutsia, Roseburia, Megamonas, Catenibacterium, Ligilactobacillus, Coprococcus, RF39* order*, Lachnospira, Megasphaera, Terrisporobacter,* and, most significantly, *Holdemanella. Holdemanella* species, associated with increased fiber consumption^41^, improved glucose tolerance^42^, and overall health^43^, and have also been suggested to be protective against colorectal cancer^44^. Taxa significantly enriched in all other cohorts compared to SAMBAR are *Bacteroides, Escherichia/Shigella, Parabacteroides, Alistipes,* Ruminococcaceae *CAG-352*, and, most significantly, *Collinsella*, which was almost absent in the SAMBAR cohort*. Collinsella* species have been associated with cancer^45^, inflammation in rheumatoid arthritis^46^, and decreased dietary fiber consumption^46,47^.

### Microbial co-abundance relationships unique to SAMBAR

The Human Microbiome Compendium showed that not only are specific taxa differentially abundant in different world regions, but also, relationships between taxa vary across regions. We used SparCC to calculate two microbial correlation networks by dividing the dataset into SAMBAR (Fig. 2D; Fig. S4) and all other regions (Fig. 2E; Fig. S5). Two notable correlations were identified as present only in SAMBAR. The first was a module of three correlated taxa: *Bifidobacterium*, *Ligilactobacillus*, and *Megasphaera. Bifidobacterium* is significantly enriched in Northern Africa and West Asia compared to SAMBAR, and is enriched in SAMBAR compared to Eastern and South-Eastern Asia, Europe and Northern America, and Sub-Saharan Africa.

Meanwhile, *Ligilactobacillus* and *Megasphaera* are both significantly enriched in SAMBAR compared to all other regions. The association of *Bifidobacterium* with dairy and whole grain consumption has long been recognized^48–50^, and recent work has suggested a relationship wherein *Megasphaera* proliferates on the metabolites produced by *Bifidobacterium*^51^. *Ligilactobacillus*, while less commonly associated with dietary factors in humans, has been used as a food additive for its probiotic effect^52,53^.

Additionally, the genus *Megamonas*, significantly enriched in SAMBAR compared with all other cohorts, formed a network including *Bacteroides* and *Ruminococcus gnavus group. Bacteroides*, significantly less abundant in SAMBAR, is associated with industrialized lifestyles in general^8,54,55^. Meanwhile, *Megamonas* and *R. gnavus* have both been associated with poor health outcomes; *Megamonas* with obesity^56^ and *R. gnavus* with Crohn’s disease and inflammation^57^. In summary, a classic urbanization marker forms a correlation network with metabolic disease-associated taxa only within SAMBAR, suggesting a potential role for these taxa in the rise of “diseases of urbanization” within South Asia.

### SAMBAR microbiomes reflect diet and lifestyle associations and heterogeneity across communities

While a broad taxonomic overview of the full SAMBAR dataset (Figs. 3A and B) compared with the compendium highlights families that are enriched in SAMBAR – Erysipelotrichaceae, Selenomonadaceae, Succinivibrioceae, and Peptostreptococcaceae – we expect microbiome heterogeneity within SAMBAR considering the geo-cultural range of our sampling. We used MaAsLin2 to perform differential abundance analysis to identify taxa enriched in each community (Fig. 3C). To select one of the communities to use as the reference, we calculated the mean relative abundance for each taxon, and identified Kolam as the community whose average composition had the smallest Aitchison distance to this mean (Table S7). The most significant community-associated genus, with the strongest effect size, was *Bacillus*. This genus was enriched (adjusted p = 9.26E-79) in the Pochury community of Nagaland in northeastern India, and in no other community in the dataset. Notably, many Pochury participants reported in the survey regular consumption of fermented legumes, specifically *axone* (Table S1). This fermented soybean product common in Nagaland has been shown to have a microbial profile dominated by *Bacillus* species^58^. We additionally observe *Ligilactobacillus* being significantly enriched (adjusted p = 0.0003) in the Spitian community of high-altitude pastoralists, who also demonstrated significant enrichment in *Bifidobacterium* and *Megasphaera* – this may reflect the regular consumption of milk, yogurt, and wheat products reported by members of this community (Table S1).

**Figure 3:**
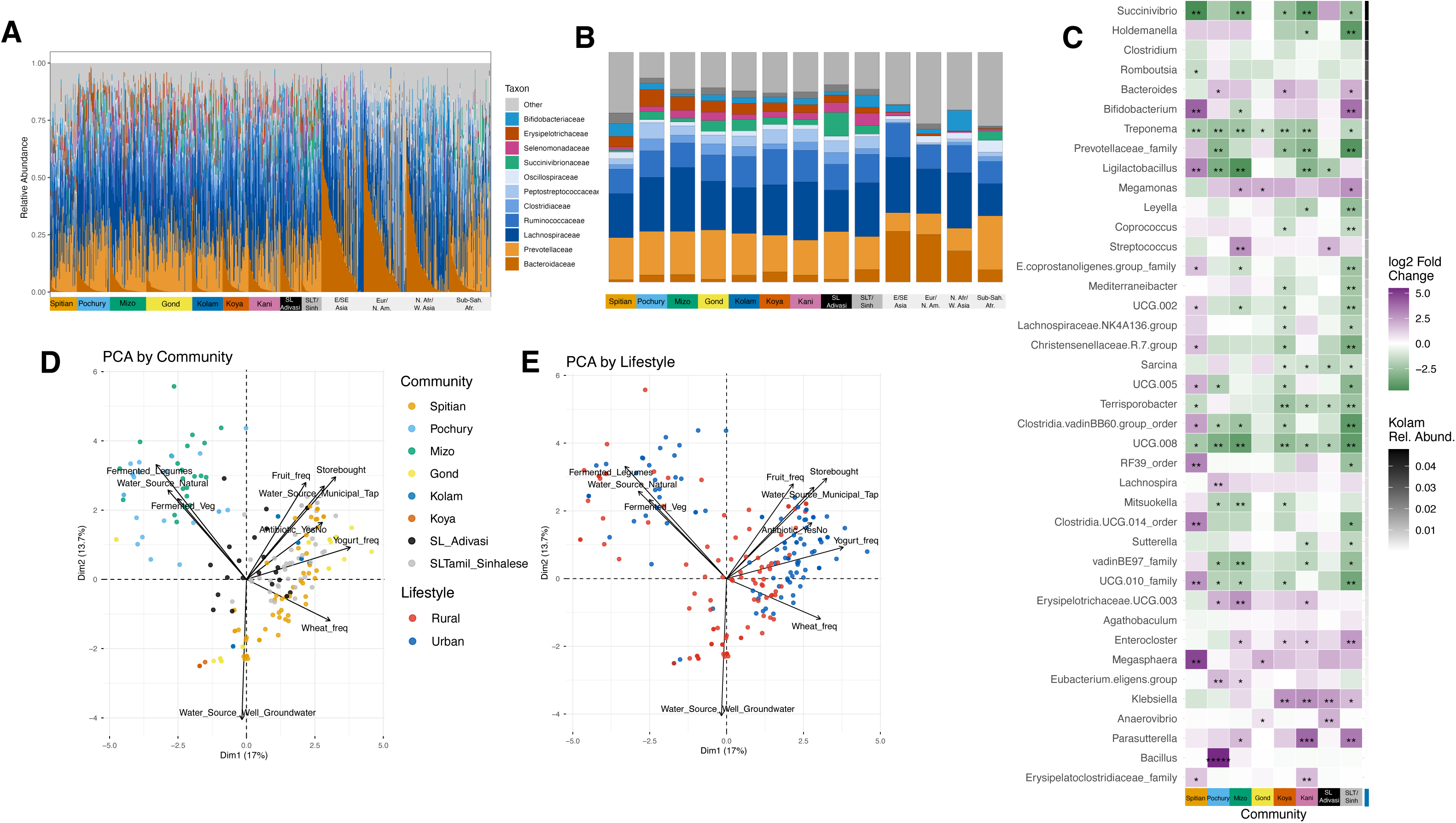
A: Stacked bar plot of all SAMBAR samples, with 100 random samples from each global cohort; taxa shown at family level. B: Stacked bar plot of taxa shown in 3A, showing the average abundance per family for each community. C: PCA of all SAMBAR individuals with complete diet and lifestyle survey data, colored by community. D: PCA of all SAMBAR individuals with complete diet and lifestyle survey data, colored by urban/rural status. E. Differentially abundant taxa between each community in SAMBAR cohort and the Kolam, controlling for lifestyle. Green taxa are more abundant in the comparison community; purple taxa are more abundant in Kolam. Asterisks reflect statistical significance as follows: adjusted p-value < 1e-50 = *****; < 1e-20 = ****; < 1e-10 = ***; < 0.001 = **; < 0.05 = *,. Taxa are ordered by relative abundance in Kolam, represented by black bars.

Beyond these three taxa, the Spitian community continues to distinguish itself from the rest of the SAMBAR cohort with significant enrichment in *RF39* order, Clostridia *UCG-014,* Christensenellaceae *R7, UCG-010,* and Oscillospiraceae *UCG-005*. Other communities demonstrated enrichment and depletion of other taxa (Fig. S6A-H); for instance, *Parasutterella* was strongly enriched in the Kani and less strongly enriched in Koya, Mizo and Sri Lankans. The documented association of *Parasutterella* with BMI and type 2 diabetes^59^ suggests that community-specific gut microbiome taxa have potential health implications. *Succinivibrio*, identified in previous studies of Indigenous microbiomes compared to urbanized^60^, was also enriched in the Sri Lankan Adivasi. Additionally, we replicated most of these results using an external reference, Sub-Saharan Africans (Fig. S6I), selected from the global panel as the most similar to SAMBAR based on PCs 1 and 2.

### Integrating microbiome and metadata shows strong effect of geography

The sampled communities spanning India and Sri Lanka share genetic and linguistic characteristics and community-specific microbiome signals, as noted above, as well as culturally specific diet and lifestyle characteristics (Table S1). The first principal component of a diet and lifestyle PCA separates the Mizo and Pochury from the rest of the cohort, distinguished by their regular consumption of fermented meat and fermented legumes, and low consumption of wheat (Fig. 3D). Also distinguished is the high-altitude Spitian community, whose wheat and yogurt consumption is markedly higher than all other cohorts (Figure 3D). However, some communities also showed separation along an urban-rural gradient (Figure 3E). For instance, the Gond, Sri Lankan, and Kolam individuals showed a significant separation between urban and rural communities, where urban communities were characterized by consumption of yogurt, wheat, store bought foods, tap water, and antibiotics. Given the heterogeneity between communities with regard to urban-rural distinction, we additionally plotted a “normalized PCA” in which each individual’s coordinates were normalized by the community-specific mean, demonstrating a more uniformly clear urban-rural separation (Figure S7). These results illustrate that the markers of lifestyle transitions from rural to urbanized lifestyles are heterogeneous across communities, and a community-specific approach would be necessary for characterizing microbial signatures of lifestyle transition.

To investigate the overall relationship between lifestyle, geography, and microbiome in SAMBAR, we built a geographic distance matrix as well as a distance matrix based on the diet and lifestyle survey and tested for correlation with the microbiome distance matrix. We found that microbiome composition is significantly correlated with geography when controlling for diet/lifestyle (partial Mantel test, p = 0.0087) but that the reverse is not true; the microbiome is not significantly correlated with lifestyle when controlling for geography (p = 0.67). We additionally implemented Spectral Correlation Analysis of Layered Evolutionary Signals (SCALES)^61^, a hierarchical clustering method, to identify factors contributing to the PC variance in microbiome composition. We observed that the most significant contributors to variance in the microbiome were features associated with geography and community membership, with all of the geography-associated variables captured in the first 12 PCs, further supporting the finding that geography is significantly correlated with the microbiome (Figure S8).

Together, our results show that diet and lifestyle play a critical role in distinguishing communities and shaping their gut microbiomes within the SAMBAR dataset. However, given these communities do not represent extremely urbanized contexts, geography and community membership are observed to have a stronger correlation with the gut microbiome composition of SAMBAR populations than lifestyle-associated dietary variables.

### Differential taxa abundance between urban and rural communities identify lifestyle and health associations

To account for the heterogeneity across communities when evaluating lifestyle transition, we calculated the differential abundance of taxa between urban and rural lifestyles, setting community membership as a random effect. While for all Indian communities, we recruited rural and urban members of the same community, the Adivasi of Sri Lanka had no urban equivalent. As such, in all lifestyle-level comparisons, the urban counterparts to the Sri Lankan Adivasi are the Sinhalese and Sri Lankan Tamil.

Overall, there were more taxa that were enriched in rural lifestyles than with urban lifestyles (Figure 4A). This is concordant with previous studies reporting that rural communities harbor microbes that can be lost during the transition to urban lifestyles^62^. The taxa with the highest effect sizes associated with rural lifestyles were *Prevotellaceae*, *Succinivibrio*, and *Holdemanella* (Figure 4A). *Prevotellaceae* and *Succinivibrio* are associated with rural lifestyles in multiple studies and are well understood to be core VANISH taxa. *Holdemanella* has been less well characterized in terms of lifestyle transition, but two studies in African populations have identified it as similarly enriched in rural communities^7,63^. Our previous results showed that *Holdemanella* is significantly enriched in the SAMBAR cohort compared to other global cohorts (Fig. 2C); its association within SAMBAR with rural lifestyles suggests that it is a potential region-specific, health-promoting taxon that is being lost as populations transition to more globalized, industrialized lifestyles.

**Figure 4:**
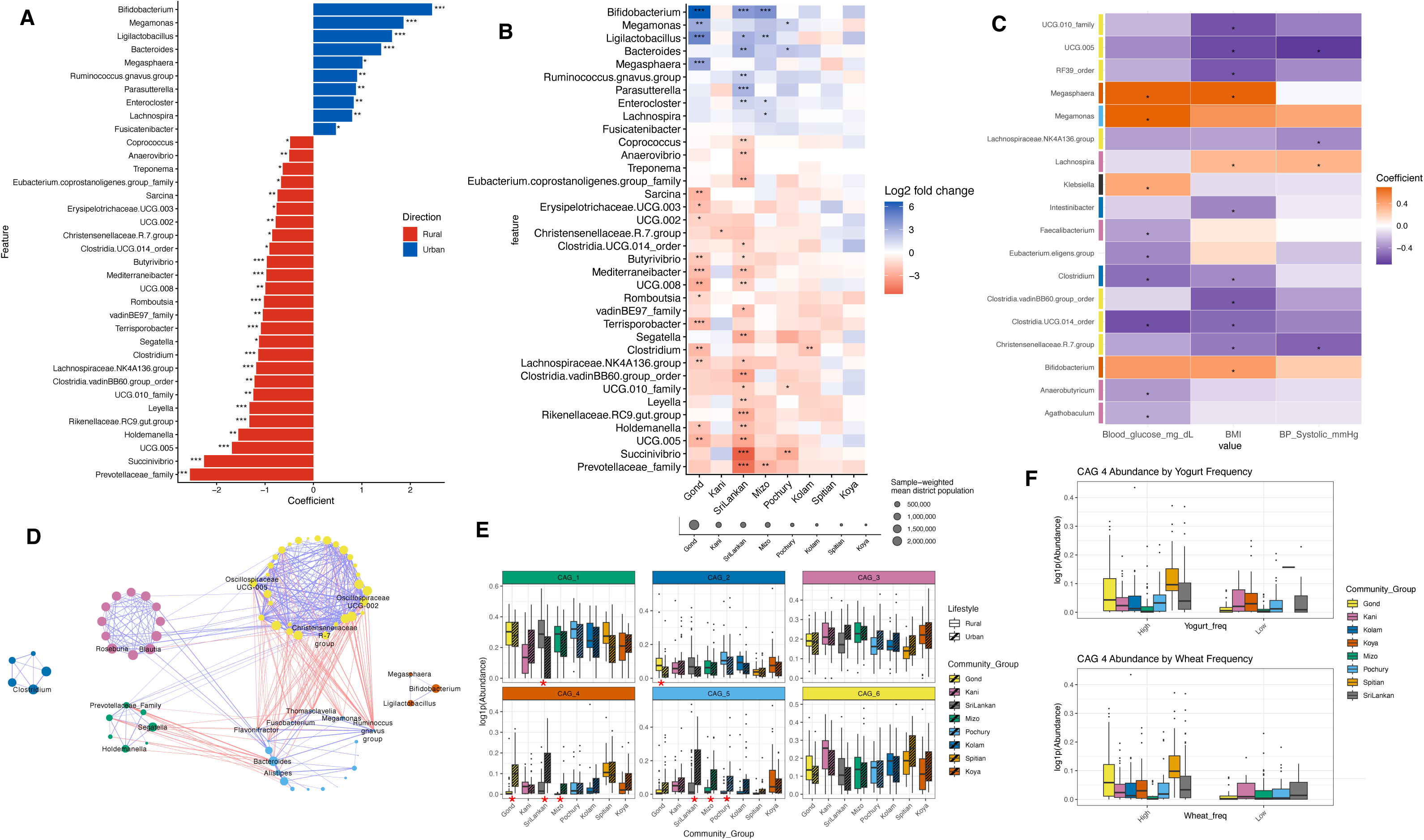
A: Differentially abundant taxa across urban versus rural lifestyles in SAMBAR, controlling for community membership. B: Differentially abundant taxa across urban and rural lifestyles within each SAMBAR community. Red taxa are more abundant in rural communities and blue in urban communities. Asterisks reflect statistical significance as follows: adjusted p-value < 1e-50 = *****; < 1e-20 = ****; < 1e-10 = ***; < 0.001 = **; < 0.05 = *. Communities are ordered from largest to smallest urban sampling location. Weighted average of urban site population size for each community is represented by grey circles. C: Heat map of taxa with significant associations with any of three phenotypes associated with metabolic disease: systolic blood pressure, blood glucose, and BMI. Orange taxa are positively correlated with a phenotype; purple taxa are inversely correlated. Bars on the left show which co-abundance group each taxon belongs to. D: Co-Abundance network showing correlated taxa within SAMBAR cohort. Top 6 most abundant CAGs are shown here; full network is shown in Fig. S13. E: Relative abundance of each CAG shown in 4C, between urban and rural counterparts for each community. Significant differences between urban and rural lifestyles are marked with red stars. F: Relative abundance of CAG 4 in “high” and “low” dairy and wheat consumers, separated by community.

On the other hand, certain taxa were associated with increased abundance in urban populations. The largest effect size was for *Bifidobacterium*, followed by *Megamonas*. These two taxa are representative of two significant trends in urbanization. We observed that *Bifidobacterium* is significantly more abundant in the urban cohorts from the Sri Lankan, Mizo, and Gond communities, compared to their rural counterparts (Figure 4B). While Koya and Kolam individuals show this effect in the same direction, the result for those two communities was not statistically significant. An overall similar effect was observed for *Ligilactobacillus* and *Megasphaera*, taxa that we identified above as being particularly enriched in the pastoralist Spitian community in concert with *Bifidobacterium*. In fact, we observed that *Megasphaera* was not only associated with urban lifestyles (Fig. 4A), but also associated with two urbanization-associated health phenotypes (Fig. 4C).

The taxon with the next greatest effect size associated with urban lifestyles, significantly in Pochury and Gond communities and trending in Mizo, was *Megamonas*. In previous studies, *Megamonas* has been shown to be positively associated with obesity, through a mechanism whereby it increases lipid absorption^56^. It has additionally been associated with HbA1c, an indicator of diabetes^64^. Of the 19 taxa associated with one of three markers of metabolic disease (BMI, blood glucose, or systolic blood pressure, after controlling for age, gender, sequencing depth, lifestyle, and community), *Megamonas* was indeed significantly positively associated with increased blood glucose (adjusted p = 0.0027), and trended towards association with increased BMI (adjusted p = 0.058) (Fig. 4C, Fig. S9). Furthermore, after controlling for age, sex, read count, and community, urban lifestyle was significantly associated with increased blood pressure (mixed model p = 0.003) and increased BMI (mixed model p = 3.54 e-05) (Fig. S10). Together, the increased prevalence of *Megamonas* in the urban cohort suggests one potential mechanism for the increase in markers of metabolic disease that may accompany urbanization. We, additionally, found that *Lachnospira*, one of the taxa significantly enriched in SAMBAR over the global reference cohorts, was also associated with urban lifestyles (Fig. 4A) and BMI and systolic blood pressure (Fig. 4C).

In contrast, several taxa were associated negatively with the markers of metabolic disease. *UCG-005* (*Oscillospiraceae*)*, Clostridium*, *Clostridia UCG-014,* and *Christensenellaceae R7 group* were each significantly negatively associated with two of the three markers, with the third trending in the same direction (Fig. 4C). Notably, this negative relationship between *Christensenellaceae* and obesity is well-documented in multiple cohorts ^65^. *Oscillospiraceae*, meanwhile, and other members of the *Oscillospira* class, have been previously associated with leanness^66^; *Oscillospiraceae UCG-005* abundance is indeed inversely related to BMI (p = 1.7 e-05) (Fig. S11).

### SAMBAR bacterial networks reflect microbiome diversity and lifestyle factors

While individual taxon associations shed light on some aspects of the microbiome, the gut microbiome is a complex ecosystem. To test the effect of urbanization on communities of taxa, we built a co-abundance group network. The resulting network showed taxa clustering into ten co-abundance groups (CAGs), with the largest six CAGs accounting for, on average, 88% of the composition of each group (Fig. 4D, Table S8; Fig. S12). Of these six, CAG 5, in light blue, was characterized by *Bacteroides*, *Parabacteroides*, *Alistipes*, and *Bilophila*, all associated with industrial diets, and particularly consumption of animal protein^67^. Indeed, abundance of CAG 5 was higher in most urban cohorts in comparison to their rural counterpart, with significantly higher abundance in urban Mizo, Pochury, and Sri Lankans (Fig. 4E, Fig. S13). In contrast, CAG 1 (green) was characterized by *Segatella*, *Leyella*, *Holdemanella,* and *Prevotellaceae*; more commonly associated with plant consumption and non-industrialized lifestyles^68^. In addition to the definitional positive correlations between taxa within CAGs, strong negative correlations (red connections) were seen, as expected, between CAGs 1 and 5 (Fig. 4E).

CAG 6 (yellow), characterized by *Oscillospiraceae, Christensenellaceae, and Eubacterium coprostanoligenes,* was the most dense network identified in this dataset. Taxa in this CAG have been associated with positive health outcomes, and in particular, lower inflammation^65,69–71^. Additionally, relative abundance of this CAG, as well as of individual taxa within the CAG, were strongly positively associated with gut microbiome diversity (Figs. S11, S14). With this in mind, a notable finding from the CAG structure is that the urbanization-associated CAG 5 was strongly negatively correlated with the diversity-associated CAG 6, as illustrated by the high density of red connections between taxa in these two CAGs. By virtue of the two CAGs connected only through negative edges, they fulfill the “two competing guilds” framework of microbial networks^72^, suggesting an oppositional relationship between the taxa associated with urbanization and the taxa associated with diversity and decreased inflammation. In particular, two CAG 5 taxa – *Megamonas* and *Ruminococcus gnavus* – were notable contributors to the negative relationship between these two CAGs. These taxa also show a strong negative relationship with diversity (Figs. S9, S15). These two taxa being significantly associated with urbanization in the full SAMBAR dataset (Fig. 4A) suggests a potential mediator for the effect of urbanization on gut microbial diversity, and consequential impacts on health, in particular inflammation.

As noted previously, the correlation network of *Bifidobacterium*, *Ligilactobacillus*, and *Megasphaera* is unique to SAMBAR compared to the four other global regions in the cohort. In Mizo, Sri Lankan, and Gond, the abundance of this CAG (the orange CAG 4) is associated with urbanization; in contrast, the pastoralist Spitian community shows significant amounts of CAG 4 in both urban and rural settings (Fig. 4E). This CAG is dominated by *Bifidobacterium*, which has been shown to be associated with whole grain diets as well as with dairy consumption^49,73–79^. In our survey of metadata, both yogurt frequency and wheat frequency were among the top features contributing to variation in the dataset in the direction of urbanization (Figs. 3C-D). As such, we hypothesized that wheat and yogurt consumption may be associated with the abundance of *Bifidobacterium*, *Ligilactobacillus*, and *Megasphaera.* We tested this by using the survey data to classify individuals as “high” and “low” consumers of wheat and yogurt and found a correlation between wheat and dairy consumption (χ^2^ test, p =0.01). To account for this confounding, we modeled taxon abundance with both dietary variables as covariates and observed that CAG 4 abundance was significantly higher in the “high” consumers of both wheat (adjusted p = 0.00004) and yogurt (adjusted p = 0.0205) (Fig. 4F). At the taxon level, we found *Bifidobacterium* to be significantly associated with both wheat (adjusted p = 6.88e-07) and yogurt (adjusted p = 0.0012). *Ligilactobacillus* was significantly associated with wheat as well (adjusted p = 9.89e-05) as was *Megasphaera* (adjusted p = 9.27e-05), but neither was associated with yogurt when controlling for wheat. As illustrated in the diet PCA (Figs. 3C-D), wheat and yogurt are both markers of urbanization in specific communities. Higher wheat consumption is significantly associated with urban lifestyles in the Gond, Kani, Mizo, and Pochury, while higher yogurt is significantly associated in the Gond, Mizo, Pochury, and Sri Lanka (χ^2^ test, Fig. S16). Notably, *Bifidobacterium* has been strongly associated with lactase nonpersistence genotypes^80–83^, and particularly with dairy consumption in lactase-nonpersistent individuals^84–86^. Given this, we screened for known lactase persistence alleles^87,88^ in SAMBAR populations or related communities drawn from previous publications, and found limited evidence for lactase persistence alleles in the majority of communities (Table S9); that is, the SAMBAR association between yogurt consumption and *Bifidobacterium* abundance is in lactase-nonpersistent individuals. Our results, therefore, suggest that the rise in *Bifidobacterium* and correlated taxa is a marker of urbanization in specific South Asian communities, and reflects the urban individuals’ increased consumption of wheat and/or yogurt.

## Discussion

A primary objective of the SAMBAR project was to build a microbiome cohort capable of elucidating the impact of ongoing lifestyle transitions on the health of traditional communities, while simultaneously leveraging anthropological genetics to represent diverse host ancestries and minimize genetic confounding within each community. To accomplish this, we employed a matched sampling design in which rural and urban participants from the same community (except in Sri Lanka) were sampled to ensure similar genetic ancestries. An additional but equally critical aim was to rectify the underrepresentation of South Asian populations, a region characterized by remarkable genetic and cultural diversity, in the worldwide gut microbiome literature. Our work increases the number of South Asian individuals in the Human Microbiome Compendium by around 10%, and the number of adult South Asians by 1500%, including the first gut microbiome data, to our knowledge, from traditional Adivasi community members in Sri Lanka. The resulting SAMBAR cohort, thus, provides insight into not only the cultural diversity of South Asian microbiomes, but also the signals of lifestyle transition within these populations. Some of these key insights were gained from harmonization with the Human Microbiome Compendium, which was possible due to comparable 16S datasets. While future whole-genome sequencing efforts would certainly allow more insight at the species and strain level, and increased power to detect immediate and emerging health impacts such as antibiotic resistance, we discuss below key results from our dataset that are nevertheless both region-specific and clinically relevant.

When comparing the SAMBAR cohort to other world regions, we identified taxa with potential health implications that are significantly enriched in South Asia, including *Erysipelotrichaceae* and *Peptostreptococcaceae* families, and specific genera including *Holdemanella*, *Romboutsia*, and *Catenibacterium*. *Holdemanella* has been associated with improved glucose tolerance and decreased inflammation^42,89^. However, both *Holdemanella* and *Catenibacterium* species have been associated with unhealthy serum lipid profiles^90^. Functional and metabolomic studies would be necessary to establish the mechanism of action of these taxa in South Asian contexts, especially how they may contribute to health outcomes in the region. *Ligilactobacillus*, another taxon at higher abundance in the SAMBAR cohort than the global comparison set, is a genus of lactic acid bacteria split from the larger *Lactobacillus* genus in 2020^91^. While *Ligilactobacillus*’s recent reclassification means that specific research into it has not been extensive, studies have evaluated members of the genus, including one isolated from a tribal Indian community^92^, as potential probiotics^52,53^. *Ligilactobacillus salivarius* has been shown to decrease colonic inflammation and intestinal damage in response to the pathogen *Salmonella* and is proposed to have a regulatory effect on the microbiota^93,94^. *Ligilactobacillus* is associated with wheat consumption within SAMBAR, though, interestingly, we do not observe an enrichment of this taxon in global regions such as Europe and North America, where diets contain significant amounts of wheat. Future studies are needed to fully understand the dietary associations and functional profiles of *Ligilactobacillus* across these regions.

One of the strongest and consistent signals differentiating SAMBAR from comparative global populations is the co-occurrence of *Ligilactobacillus* with *Bifidobacterium and Megasphaera.* In SAMBAR, *Bifidobacterium* is associated with consumption of both wheat and dairy, specifically yogurt. *Bifidobacterium* has long been associated with dairy consumption, especially in individuals with lactase non-persistence^86^, and has even been used as a probiotic. Supplementation of *Bifidobacterium*-enriched yogurt has been shown to mitigate maldigestion in lactase non-persistent individuals in multiple studies^95–97^. Given the SAMBAR cohort is largely expected to lack genetic adaptations for lactase persistence, our findings point to non-genetic mediators, including the gut microbiome and/or yogurt fermentation, as potential adaptive strategies in these South Asian communities, paralleling observations in Central Asian pastoralists^98^. *Megasphaera*, on the other hand, has been shown to utilize metabolites produced by *Bifidobacterium*^51^, possibly explaining the correlation between these two taxa. *Megasphaera* itself has been used as a probiotic in dairy cows^99,100^, and has been suggested as a probiotic for humans ^99^. It is therefore worth investigating whether *Megasphaera*’s presence in the SAMBAR cohort reflects any bovid husbandry practices that increase either human or animal wellbeing.

Additionally enriched in the SAMBAR cohort was *Megamonas*, whose health impacts have been well characterized. Importantly, within the SAMBAR cohort, *Megamonas* has one of the strongest associations with urban lifestyles. In South Asia, as in published works, *Megamonas* has a clear association with obesity, and one mechanism for this has already been elucidated^56^. Additionally, *Megamonas* shows a strong negative correlation with microbiome diversity in SAMBAR. The enrichment of *Megamonas* in South Asian urban populations suggests a potential contributor to the increased rates of metabolic disease and obesity in South Asia, and highlights the need for more population-specific studies, including integration with metabolomic profiles. Also part of the co-abundance group with *Megamonas* and urbanization-associated taxa are *Ruminococcus gnavus group*, *Fusobacterium*, and *Flavonifractor.* All three of these taxa have been associated negatively with health; *R. gnavus* with Crohn’s disease^57^, *Fusobacterium* with colorectal cancer^101^, and *Flavonifractor* with colorectal cancer specifically in an Indian cohort^102^. All taxa showed strong negative correlations with the anti-inflammation-associated CAG 6 in SAMBAR.

As the most comprehensive South Asian gut microbiome dataset so far, the SAMBAR cohort also highlights the strong geographic structure and heterogeneity in gut microbiomes seen across communities and lifestyles, underscoring the need for a local/region-specific approach to sampling of more diverse populations. For example, we observe stronger differences between urban and rural cohorts in the Gond and Sri Lankans, as evidenced by the increased number of significant taxa in these communities (Fig. 4B). We additionally see that the *Bifidobacterium* and *Megamonas* associations with urbanization are not homogeneously seen in all communities (Fig. 4B). This heterogeneity reflects both the cultural and dietary diversity in South Asia, but also sheds critical light on the nature and rate of lifestyle transitions that are likely more nuanced than the urban-rural dichotomy^103,104^. Most of the sampled rural communities within India maintain some, though varied, degree of access to the industrialized world (see ‘Community Descriptions of Diet and Lifestyle Shifts’ in Methods). Similarly, although none of the sampled Indian urban cohorts are from any of the top ten largest cities in India, the Gond urban migrants originate from the largest urban center represented in our cohort. Together with the Sri Lankan urban donors (Sinhalese and Sri Lankan Tamil), who reside in some of the largest cities in the country such as Colombo and Kandy, these communities are therefore expected to exhibit the strongest urban-rural contrasts, assuming urban population size is a reasonable proxy for the degree of “urbanness” (Fig. 4B). In contrast, the Koya exhibit among the least pronounced urban-rural difference as the Koya urban migrants preferentially move to (and were, thus, sampled from) smaller urban centers or even villages rather than large cities (Fig. 4B). One notable exception to this trend is the Kani, who despite the large size of the urban center, show minimal differences between the urban and rural microbiomes (Fig. 4B); this likely reflects greater access to industrialized foods in rural community members. While the urban-rural designation is practical, we recognize that the urbanization signal in the microbiome data, in practice, is likely reflecting a mix of both individual-level choices and, at a broader scale, access and infrastructural options available to the urban migrants (e.g. whether they move to cities versus towns or villages, readily available processed foods, the extent to which they actively change their diet and lifestyle upon migrating).

Overall, our research characterizing the microbiome of this vastly underrepresented region highlights unique signatures of the diverse populations and lifestyles, and emphasizes the importance of future study in this region. Importantly, with increased regional studies, close collaboration with local leaders, community members, and researchers with existing relationships is necessary to continue building this dataset in an ethical and anthropologically-informed manner^105^.

## Materials and Methods

### Outreach and Participant Recruitment

We conducted an anthropologically informed survey of select Indian and Sri Lankan populations in collaboration with local scientists, anthropologists, archaeologists, ethnobiologists, and community members with longstanding roots in each community. Within India, populations generally span a genetic cline from north to south shaped by differential affinities to regional ancient hunter-gatherers and various West Eurasians, while many Northeast Indian populations are genetically more similar to East Asian populations^106–110^. Genetic affinities generally, with notable exceptions like the Dravidian-speaking Gond, mirror linguistic families, with geographically and genetically close populations often speaking languages within the same broad language family^14^. We designed our sampling strategy to span this vast genetic and cultural gradient within India, recruiting from communities across a broad geographic and linguistic range. In Sri Lanka, the Adivasi are an Indigenous community who are anthropologically proposed to be descendants of ancient hunter-gatherers and are reported to have slightly higher genetic affinity to ancient hunter-gatherers than the more recently arrived Sri Lankan Tamil and Sinhalese^33^. Many members of the Adivasi community continue to use hunter-gatherer subsistence strategies today, although they are subject to relocation and habitat fragmentation. Our final cohort consisted of individuals from the following communities: Gond, Spitian, Kani, Kolam, Koya, Mizo, Pochury (India) and Adivasi, Sinhalese, and Sri Lankan Tamil (Sri Lanka). Our metadata (Table S1) and sampling map (Fig. 1) aggregate urban and rural donors for each community at the district level to maintain anonymity, given some settlements are small.

Members of the research team visited rural communities across India and Sri Lanka with materials explaining the project goals and methods, translated into each local language. We first received consent from state and local government bodies and local community leaders or representatives to include their village in our sampling, after which we held public meetings in each location to explain the project to all interested community members. After these meetings, each eligible participant engaged in our informed consent process to be enrolled in the study. Individual participants within each community were recruited based on additional criteria: age between 18 and 60, no antibiotic use within the last 3 months, and not a first-degree relative of any individual already enrolled. We aimed to enroll roughly equal numbers of men and women. The project was subject to the Institutional Review Board at the University of Chicago, USA (IRB19-1761); the Institutional Ethical Committee at the Birbal Sahni Institute of Palaeosciences, India (BSIP/Ethical Approval/2021/Letter-1); and the Ethics Review Committee at the University of Colombo, Sri Lanka (EC-17-147). In-person return of results to the participating communities is planned concurrently with the peer review process. As with the initial outreach, this will be done via presentations conducted with the help of community contacts and translators, additionally accompanied by written materials/visual aids translated into local languages.

### Community Descriptions of Diet and Lifestyle Shifts

This section summarizes key dietary components and signs of lifestyle shifts in each of the sampled communities, deriving primarily from observations and unstructured interviews by the research team and community knowledge, as well as available ethnographic records.

#### Gond

The subsistence strategy of the sampled Gond community is traditionally characterized by resilience and sustainable utilization and dependence on natural resources. Millets, including varieties such as kodo, kutki, ragi, and jowar, form the core staple diet and are consumed in the form of rotis, steamed cakes, and *pej/Ambil*, a cooling millet gruel widely consumed during summers. Locally grown upland rice is also used in daily meals as well as in fermented foods. Alongside cultivated grains, forest produce plays a crucial role, providing essential micronutrients and acting as a buffer against seasonal scarcity. Wild edibles such as mahua flowers and seeds, tendu, tamarind, bamboo shoots, mushrooms like *rugda and putu*, and diverse tubers constitute a substantial component of traditional diets and knowledge systems. Local poultry, goat meat, freshwater fish, and wild boar are also consumed seasonally. A notable cultural food is *chapda chutney*, made from red weaver ants, valued for its tangy flavour and medicinal properties. Edible insects, including termites and silkworm larvae, supplement nutrition especially during monsoons. Along with leafy green vegetables, traditional foods such as *bafauri* (steamed gram-flour dumplings) and *farra* (rice-flour dumplings) constitute low-oil, nutrient-dense culinary preferences. Fermentation is central to Bastar’s food culture, with *handia* (rice beer), mahua-based, and palm-based (*tadi*, *gorga*) brews being integral to ritual and social life. Fermented bamboo shoots and rice-based preparations demonstrate a sophisticated understanding of preservation technologies suited to a monsoon-adapted environment. Rural community members also ferment milk to prepare curd/yogurt and *kadhi* (buttermilk) using excess cattle milk.

With seasonal migrations and improved access to cities, rural households are beginning to gain exposure to urban dietary habits. Migrant workers returning from cities frequently introduce foods such as packaged *masala* (spices), instant noodles, frozen items, and restaurant-cooked foods, which are gradually diffusing into the wider community. Additionally, through ‘mid-day meals’ and government-sponsored grocery provisions, or *rations*, are other mechanisms of introduction of non-local food items such as processed rice, lentil, sugar and palm oil.

#### Spitian

Yak milk forms a core component of the traditional diet of the high-altitude dwelling Spitians. While there is no significant seasonal variation in milk consumption, cow milk is increasingly consumed nowadays due to a decline in the local yak population. Zomu, a crossbreed of yak and cow, is another local source of milk. Milk is consumed both raw, especially in tea, and in the form of curd and cheese. Spitians traditionally prepare a fermented cheese, called *chhurpe*, which is eaten in its dried form throughout the year. In addition to dairy, barley is a key staple and eaten in different forms. In fact, many urban migrants take *sattu*, powdered barley that is often reconstituted with yogurt, from their villages and consume it when they visit their homes. Despite an effort to retain this traditional dietary component, grains like wheat are becoming increasingly common among urban migrants.

#### Kani

The Kani tribal community of the southern Western Ghats, inhabiting the forest regions of Thiruvananthapuram and Kollam districts in Kerala and the adjoining areas of Tamil Nadu, maintains a distinctive forest-based food system centred on rice, millets, tapioca, elephant yam, taro root, wild mushrooms, and more than 50 wild edible plant species. These include foraged tubers (*Dioscorea* spp.), leafy greens, seasonal fruits (*Artocarpus hirsutus*, *Baccaurea courtallensis*, *Syzygium* spp., and wild mangoes), and honey. Their subsistence strategy combines foraging and small-scale cultivation. Traditionally, animal protein has been obtained through hunting, including wild boar and small animals. Rural Kani households use traditional food-processing methods, with wild edibles typically cooked as accompaniments to rice using salt, chilli, coconut, and onion.

However, improved road networks and market access have increasingly exposed these communities to packaged snacks, wheat-based products, and sugar-sweetened beverages. Approximately half of the Kani population now lives in urban areas (township-based) and predominantly relies on urbanized foods. Even for those still living in rural settings, the distinction between Kani rural and urban populations is minimal. Although they are geographically located in rural areas, they have access to facilities and amenities comparable to those available in urban settings. Urbanization is marked by substantially higher consumption of refined wheat and rice products, as well as commercially prepared foods, including fish and meat. This reflects selective dietary transitions toward convenience foods facilitated by market integration, although community-specific ethnographic documentation is limited. Notably, curd and yogurt consumption represents a new dietary practice rather than a traditional cultural element, as dairy products have historically been absent from Kani food culture because of the limited availability of dairy livestock in forest habitats. Traditionally, protein sources included forest foods such as tubers and legumes, along with hunted wild animals. Today, both rural (forest-dwelling) and urban (township-based) Kani communities receive food supplies through the Government’s Public Distribution System (PDS), which includes rice, wheat and pulses.

#### Kolam

Traditionally, the Kolam community followed a small-scale hunting and foraging lifestyle. The core diet consisted of *jonna* (jowar or sorghum), an agroforestry-based crop, typically consumed in the form of *jonna ambali* (a malt-based preparation consisting of *jonna* and *mokka jonna* (maize)). The *vippa chettu* or *mahua* (*Madhuca longifolia*) was central to their diet, with flowers consumed seasonally and seeds used for oil extraction for cooking. This was obtained through foraging and supplemented with natural honey and seasonal wild fruits. Historically, households also prepared curd and buttermilk from cattle milk, but consumption of these dairy items is infrequent in rural areas these days.

Today, rural community members have growing access to processed foods and government sponsored programs (primarily rice and sugar). On the other hand, migrant Kolam individuals in towns and cities have adopted urbanized diets to a large extent. The present-day staple diet of the Kolam community consists primarily of rice obtained through the Public Distribution System and tur dal (split pigeon peas), typically prepared with red or green chillies, tomato, and seasoned with cumin seeds, onion, curry leaves, and palmolein oil. Both rural and urban Kolam households generally follow a twice-daily meal pattern. While tur dal is cultivated locally, most other food items are purchased from nearby markets. The community consumes market-sourced vegetables such as cabbage, carrots, and beans, alongside animal-based foods including chicken, eggs, and fish, and occasionally wild pork obtained through hunting. Consumption of animal products varies by economic status, occurring once or twice a week for most households, though families that continue to practice hunting may consume meat more frequently.

#### Koya

The historic dietary traditions of the Koya community reflect a subsistence pattern closely aligned with agrarian practices and utilization of the local forest ecology and pastoral resources. Their staple food system was centred on *jonna gataka*, a thick porridge prepared from *jonna* (jowar or sorghum) and consumed with *dal* (lentil) and supplemented with ghee. Equally significant was *jonna ambali*. Another core component of their dietary regime was *ragi sankati*, a finger-millet-based dough commonly eaten with *kaaram* (chili powder) and ghee, indicating a strong reliance on millet cultivation and consumption. Protein sources were traditionally diverse, comprising *naatu kodi* (country chicken), beef, and a range of seasonal forest products.Occasional hunting was also part of their subsistence activities, providing meat such as wild boar and deer. Dairy, particularly curd, contributed additional nutritional value. Culinary practices predominantly employed locally sourced oils, including *ippa* (*mahua*) oil and groundnut oil.

In contemporary contexts, however, the community’s food traditions have undergone marked transformations. With expanding market integration, government rations, increased mobility, and penetration of packaged and commercial foods, there is a growing shift towards processed staples such as rice, snacks, instant noodles, soft drinks, and fast food items such as fried chicken. The proliferation of digital platforms has further enabled the consumption of online-ordered and ready-made foods, reflecting broader socio-economic changes and signalling a gradual departure from traditional millet-based dietary practices.

#### Naga Pochury

A variety of rice preparations, with processed vegetables (smoked, sun dried, or fermented) and meat (both domesticated and wild and/or worms such as woodworms) form the core diet of the Naga. Diverse cooking styles like steaming, boiling, and roasting, and the usage of different types of vessels (e.g. ceramic pottery or bamboo) introduce flavor variations in their daily diet. Rice is usually cultivated, while vegetables are either foraged or locally grown. The inherent diversity in the Pochury diet is underscored by different sampled villages having their unique cooking traditions; examples of foods specific to some of the villages include *xetri* (a sour wild leaf mixed with taro corm, Naga *dal*/lentil, salt, and small spicy chili), *phzuti* (taro corm, job’s tears millet, small spicy chili, and salt), *xauseum* (yam mixed with salt), *axone*/*khroshwü* (fermented soyabean), *tüxhüli* (sour wild leaf cooked with local lentils, fermented soya, dry king chilli, and salt), wulishi (wild fern cooked with taro or local lentils, ginger, chilli, salt, and fermented soya), *küdji* (pounded black sesame cooked with smoked pork/dry buffalo skin/snails/dry vegetables like taro stem, long beans, bamboo shoots, mushroom etc. and seasoned with Vietnamese balm, fermented soya, salt, and dry king chilli), *küüzü* (perilla seeds pounded with cooked sticky rice and salt or with dry vegetables and smoked meats), *litümüsü* (gravy soup cooked with fermented soya, salt, chilli, and seasoned with aromatic plants), and *xha*, *picha*, *shapisha*, and *süshapisha* (starchy root vegetables cooked with brine and eaten as breakfast or lunch along with tea and chutney). Dairy does not constitute a traditional food source for the Naga.

While there is widespread consumption of urban foods in urban centers and a recent uptake of urban foods among rural youth, the older generation in rural settings does not consume ultra-processed foods frequently. Examples of processed and non-local foods consumed daily may include biscuits/shortbreads, milk powder, tea leaves, sugar, pulses (lentils), cooking oil, wheat (flour), sweets, chocolates, chips, and vegetables that have a longer shelf life but not grown in the area (e.g. garlic, onions). These foods are today available in local village shops or via a family member visiting urban areas. Curd and yogurt are also recent introductions, though consumed occasionally. Milk, in its raw form or as milk powder, may be consumed daily by some individuals in the urban area, primarily in tea. *Paneer* (cottage cheese) is, again, a very recent introduction and is not a popular item cooked in a Naga kitchen. Butter may also be consumed in urban areas (and some rural areas based on availability). A small amount of dairy may be indirectly consumed through baked items.

#### Sri Lankan Adivasi

Traditionally, Adivasi communities practiced a subsistence lifestyle based on hunting and gathering. Meat and fish were commonly prepared by direct roasting over wood fires, covering with hot ashes, smoking, or drying on wooden racks. Surplus meat from hunts was sun-dried or smoked to ensure preservation during the rainy season. Protein sources also included perume (a sausage-like preparation) and other meat products such as venison, monitor lizard tail, and boneless game meat. Honey harvesting from a variety of forest insects, undertaken as a collective activity, was practiced both for direct consumption and for use in meat preservation. In recent decades, however, conservation policies and wildlife protection laws have increasingly restricted hunting practices, thereby limiting traditional food sources.

Plant-based components of the Adivasi diet have historically included forest-derived tubers and yams, primarily species of Dioscorea and, less frequently, plants from the family *Araceae*. These were complemented by locally cultivated cereals such as rice, finger millet (*Eleusine coracana*), and maize (*Zea mays*), which were processed into flour and prepared as roti (unleavened flatbread) or thalapa (a thick boiled flour paste), typically consumed with ānama (smoked meat cooked with gravy). When available, cereal flours were supplemented with cycad (*Cycas circinalis*) seed flour or dried and ground Bassia longifolia flowers to enhance roti and thalapa preparations.

The regular diet also incorporated a wide range of wild herbs, leafy vegetables, and unripe fruits of gourds and melons, many of which possess medicinal and therapeutic properties. Commonly consumed leafy greens included *Cassia tora*, *Ipomoea cymosa*, and *Memecyclon umbellatum*. Additionally, ripe wild fruits and berries-such as *Mangifera zeylanica, Nephelium longana, Hemicyclia sepiaria, Manilkara hexandra, Terminalia bellirica*, and *Dialium ovoideum-*along with wild mushrooms, formed an integral part of the diet. Condiments and spices, including cumin, coriander, green chilli, and various herbs, were used in the preparation of curries and traditional dishes such as kurakkal, deep-fried balls made from a mixture of meat or fish and roasted rice flour^111,112^.

#### Mizo

Traditionally, the Mizos lived a subsistence life, depending primarily on foods such as maize, millet, and yam. Subsequent migrations and settlement at new locations in the 16th century coincided with the beginning of rice cultivation among the various tribes, rapidly becoming the most dominant food and aspect of economic, social, and cultural life of the Mizo. Until British colonization, the Mizo population consisted of agriculturists. Rice constituted the staple food, and livelihood was sustained primarily through paddy cultivation. The significance of rice to the Mizo was such that it was generally used as tribute to the community chiefs. Non-glutinous rice was cultivated and consumed universally as the principal staple food, while glutinous rice was primarily used for brewing rice beer and for preparing rice flour, which formed the base of the popular snack *chhangban* (sticky rice flour wrapped in banana leaves prepared locally within each household). Leftover boiled rice was commonly consumed between meals, especially by children, without any accompaniment. Children would take a handful of boiled rice, known as *chawtlang*, and eat it while playing in village streets. For a Mizo child, *Chawtlang* represented the most cherished and satisfying form of snack^113,114^. In times of poor rice harvests or famine, the Mizos depended on alternative food sources such as arum, pumpkin and its leaves, millet, maize, and sweet potatoes/yams. Apart from *chhangban*, snacks largely consisted of wild fruits. There is little evidence of sweet foods such as sugar or jaggery in the pre-colonial period, despite the presence of sugarcane plants in the hills. Similarly, there is no evidence of milk production or wheat consumption in the historical Mizo cuisine, with dairy and wheat consumption likely introduced in the colonial period.

Today, there is no village in Mizoram that has no access to urban food, whether processed or not.

### Sample Collection

Each participant provided a fecal sample using the OMNIgene•GUT kit (OMR-200, DNAGenotek). Each participant answered a survey about their daily lifestyle and diet, as well as about their basic demographic information. Members of the research team collected basic medical and anthropometric measurements: height, weight, waist and hip circumference, blood pressure, and blood glucose. For those participants who consented to a finger-stick blood glucose measurement, we recorded whether they had been fasting for 8 hours before the test. These lifestyle,diet, demographic, and phenotypic attributes and donor responses can be found in Table S1.

### Sample Processing

Fecal kits were shaken vigorously to homogenize samples upon receipt. Samples were transported at ambient temperature to the Birbal Sahni Institute of Paleosciences, Lucknow, India, where they were stored at –80°C until processing. Samples were thawed before DNA extraction. DNA was extracted according to the manufacturer instructions for the DNEasy PowerSoil Pro kit (Qiagen), with two modifications:

1. 250 µL of sample was transferred from the OMNIgene•GUT tube to the PowerBead Pro tube, as specified by the “OMNIgene®•GUT microbial DNA purification protocol using QIAGEN® QIAamp® PowerFecal® Pro DNA Kit.”
2. Addition of an incubation step at 65°C for 10 minutes before the bead-beating step, as recommended in the PowerSoil Pro protocol for difficult-to-lyse samples.

Samples were homogenized using a Vortex-Genie 2 (Scientific Industries) with Vortex Adapter (Qiagen). Samples were vortexed in batches of 24, so vortexing time was increased to 20 minutes per manufacturer recommendations. Samples from all communities studied were randomized to batches of 48 for extraction to minimize technical batch effects. Extraction blanks of distilled water were processed through each batch of samples. Additionally, one aliquot from the ZymoBIOMICS Microbial Community Standard (Zymo Research) was processed as part of each extraction batch as a positive control to ensure consistency across batches.

### Library Preparation and Sequencing

The V4-V5 region within the 16S ribosomal RNA gene was amplified using universal bacterial primers – 563F (5’-nnnnnnnn-NNNNNNNNNNNN-AYTGGGYDTAAA-GNG-3’) and 926R (5’-nnnnnnnn-NNNNNNNNNNNN-CCGTCAATTYHT-TTRAGT-3’), where ‘N’ represents the barcodes and ‘n’ are additional nucleotides added to offset primer sequencing. PCR conditions included initial denaturation at 94°C for 3 min followed by denaturation, annealing and extension at 94°C/15 s, 51°C/30 s and 72°C/1 min.

Final extension was performed for 5 min. The approximately 360 bp amplicons were subsequently purified using a magnetic bead-based size selection. Purified amplicons were quantified using a QubitFlex Fluorometer (Invitrogen) and pooled at an equimolar concentration before Illumina compatible Combinatorial Dual Index (CDI) adapters were ligated using the QIAseq 1-step amplicon library kit (Qiagen). The completed libraries were sequenced on an Illumina MiSeq platform to generate 2×250bp paired-end reads.

### Bioinformatic Processing

We used dada2 (v1.18.0) for processing MiSeq 16S rRNA gene amplicon sequencing reads with minor modifications in R (v4.2.2). Specifically, reads were first trimmed at 190 bp for both forward and reverse reads to remove low quality nucleotides. Chimeras were detected and removed using the default consensus method in the dada2 pipeline. Then, ASVs with length between 320 bp and 365 bp were kept and deemed as high quality ASVs. Taxonomy of the resultant ASVs were assigned to the genus level using the SILVA v138.2 classifier as formatted for DADA2 by Abdill et al., with a minimum bootstrap confidence score of 80^16^.

The SAMBAR cohort samples were filtered to a minimum 5000 reads in concordance with the Earth Microbiome Project^115^, after which 575 samples remained. Filtering and downstream processing was done using phyloseq^116^.

### Compendium Integration

The Human Microbiome Compendium (v1.1.1) was downloaded and filtered to include only projects that, like the SAMBAR project, matched the following criteria: 1) stool samples only, 2) inclusion of adults only, 3) a bead-beating step in processing, and 4) sequencing of the V4 amplicon. The compendium was then filtered further to eliminate any projects including critically ill individuals, and any projects on gastrointestinal diseases: gastric and colorectal cancer, cholera, and *H. pylori*.

After these filters, four world regions had more than 575 individuals remaining in the compendium: Sub-Saharan Africa, Europe and Northern America, Northern Africa and Western Asia, and Eastern and South-Eastern Asia. To form the global reference panel against which to compare SAMBAR, 575 individuals were randomly selected from each of these regions after filtering.

One important limitation to this approach is that the SAMBAR samples sequence the V4-V5 amplicon, which was underrepresented in the Compendium and nonexistent after the above mentioned filtering steps. As such, we were unable to control for amplicon choice when comparing SAMBAR and non-SAMBAR cohorts. We attempted to minimize any potential bias by including only amplicons overlapping the V4 region (V3-V4 and V4) from the Compendium.

### Principal component analysis by region

The principal components analysis was Aitchison PCA; that is, the data was centered-log-ratio transformed after adding a pseudo-count of half the minimum relative abundance to all values (compositions::clr). Then, the principal components matrix was calculated between the clr-transformed individuals (stats::prcomp). Final PCA plots were made using microViz::ord_plot.

### Differential abundance analysis by region

We used MaAsLin2 to identify genera that are differentially abundant between SAMBAR and at least one of the other 4 world regions. We first identified all genera with mean relative abundance of at least 0.5% and prevalence of at least 1% in any of the five regions, as described in the Compendium paper – a list of 69 genera. The OTU tables were aggregated so that all genera not in the 69 for analysis were summed as “OTHER”, and MaAsLin2 was run with the following parameters: normalization = “TSS”, transform=”LOG”, analysis method = “LM”, correction method = “BH”. Region was set as the fixed effect for comparison, with the reference region being SAMBAR. The top 40 most significant taxa by average adjusted p-value across the 4 regions were plotted, with the full 68-taxon results in Supplementary Figure 4.

### Taxon co-abundance groups: by region

To find correlations between genera in SAMBAR and non-SAMBAR cohorts, we used SparCC^117^. The combined dataset was filtered to only genera with at least a 0.5% mean relative abundance in at least one of the five regions. SparCC was run on the filtered dataset through the SpiecEasi^118^ package, with 0.3 as the threshold for correlation. The dataset was then split into SAMBAR and non-SAMBAR cohorts, and the same taxa were used to build SparCC networks within each cohort. We used igraph to combine edges from both the SAMBAR and non-SAMBAR networks into a “main” graph, and edges in this graph were annotated with their presence in the SAMBAR graph, non-SAMBAR graph, both, or neither. A subset of this graph is in Figure 3D and E, and the full “main” graph is in Supplementary Figure 5 and 6.

Bar plots: To plot taxonomic composition at the family level for each individual, we used the entire SAMBAR dataset, and a subset of 100 randomly selected individuals from each of the other four non-SAMBAR regions. For the aggregated plot, abundance values were summed using the phyloseq::merge_samples function.

### Differential abundance by community

MaAsLin2 was used to identify taxa associated with specific communities.

The mean taxon abundance for each taxon for the full SAMBAR cohort was calculated, and mean values were also calculated for each community. The Aitchison distance was calculated between each community and the mean, identifying Kolam as the group closest to the “SAMBAR mean” taxonomic profile.

We first identified all genera with mean relative abundance of at least 0.5% and prevalence of at least 1% in any of the 10 regions to produce the taxon set of 63 taxa for differential abundance analysis. MaAsLin2 was run with the following parameters: normalization = “TSS”, transform=”LOG”, analysis method = “LM”, correction method = “BH”. Community was set as the fixed effect for comparison, with the reference being Kolam. Additional fixed effects were age, gender, number of sequencing reads, and lifestyle (urban/rural).

### Principal Components Analysis: survey data

Qualitative survey data was encoded on a numeric scale of 0-3, as described by Jha et al^9^. Binary variables were encoded as 0/3, while frequency variables were encoded on a scale from never = 0 to multiple times a week = 3. Storebought snacks and drinks were grouped into a variable called “Storebought” whose value per individual was their maximum value for either category. Red and white meat were similarly grouped together, as were milk and other non-yogurt dairy. All individuals with complete survey data (337) were included in the principal component analysis. FactoMiner was used to generate the PCA, and Factoextra was used for the plot. The top 10 survey variables contributing to variation are shown on the biplot.

### Lifestyle and Geography Correlation with Microbiome

In order to test the correlation of microbiome composition with geography and lifestyle, we calculated the pairwise distance matrices for each community, following Febinia et al^119^. Data was aggregated by taking the mean for each community, and pairwise distances were calculated between all communities.

Microbiome composition was averaged by community, and the microbiome distance matrix between communities was calculated using pairwise Aitchison distances (Euclidean distance on clr-transformed data). For anonymity, geographic coordinates for each individual’s location were replaced with the mean location of all individuals from the same district. Community latitude and longitude means were calculated as the mean of the district-aggregated coordinates. Great-circle geographic distances were calculated from the mean location of each community.

A chi-square distance matrix was calculated for the diet survey data, after the aforementioned numerical transformation. We then ran the following partial Mantel tests, as described by Febinia et al: “(microbiome divergence ∼ geographic distance | lifestyle distance)” and “(microbiome divergence ∼ lifestyle distance | geographic distance)”.

### Differential Abundance between urban and rural

We used MaAsLin2 to calculate differential abundance between urban and rural groups. The candidate taxa were selected following the same protocol as Figure 3: “Differential abundance by community”, and all other taxa were aggregated. MaAsLin2 was run with the following parameters: normalization = “TSS”, transform=”LOG”, analysis method = “LM”, correction method = “BH”.

For the SAMBAR dataset, age, sex, sequencing depth, and lifestyle were set as fixed effects, and community membership as a random effect. Significant taxa after BH correction were plotted in the diverging bar plot. For each individual community, age, sex, sequencing depth, and lifestyle were set as fixed effects.

### SCALES

We used the SCALES method^61^ to assess how information related to specific metadata are distributed across principal components (PCs) based on ASV abundance. To run this analysis, we first converted continuous metadata variables (e.g. Age) to discrete bins. Samples missing more than 25% of metadata variables were removed from the metadata and from the counts. ASVs found in fewer than 5 samples were also filtered from the counts. We then calculated the singular value decomposition (SVD) of the count matrix. In the SCALES approach, each pair of samples is represented by one vector describing the correlation between their principal components and by a second vector describing the correlation between their metadata. These values are then used to estimate mutual information (MI) between metadata and principal components across all samples and variables and can reveal a range of PCs that captures information about each variable. In Figure S8, we identified the set of PCs that explained the first 50% of MI for each metadata variable and then sorted the metadata by the median PC where their information is found.

### Taxon – Phenotype Associations

We used MaAsLin2 to model the relationship between genus abundance and three continuous phenotypes. The candidate taxa were selected following the same protocol as Figure 3: “Differential abundance by community”, and all other taxa were aggregated. MaAsLin2 was run with the following parameters: normalization = “TSS”, transform=”LOG”, analysis method = “LM”, correction method = “BH”.

For each phenotype of interest, the phenotype was set as a fixed effect along with age, sex, sequencing depth, and lifestyle. Community membership was set as a random effect. For blood glucose, an additional fixed effect was a binary variable indicating whether the participant had fasted for 8 hours before providing the blood sample. MaAsLin2 outputs from each of the phenotypes were combined and p-values were BH-corrected in the full three-phenotype dataset. Genera with any significant association with at least one of the three phenotypes were visualized in Figure 4C.

### Co-Abundance Groups

We used SparCC to calculate correlations between taxa. As before, the combined SAMBAR dataset was filtered to only genera with at least a 0.5% mean relative abundance in at least one of the five regions, and 0.3 was set as the minimum correlation to be included in the final correlation matrix. After building the graph of positive correlations only, the Girvan-Newman clustering algorithm was used to assign taxa to co-abundance groups. CAGs were then visualized on the graph of positive and negative correlations to examine the antagonistic relationships between CAGs. Taxa abundances were summed according to each taxon’s CAG membership and relative abundance of each CAG was plotted.

### Association of Taxa with Diet Variables

Food frequency data was encoded on a 0:2 scale as described previously. We ran MaAsLin2 on the taxon set of 69 taxa meeting the criteria for abundance and prevalence per region described above, using the model: Taxon ∼ Wheat_frequency + Yogurt_frequency + Age + Gender + Number of reads + (1 | Community).

### Association of Diet Variables with Lifestyle

For the binary analysis, “high” consumers were 1 and 2 (monthly to daily consumption), while “low” consumers were 0 (never to rarely). We ran a chi-square test for each community to test whether yogurt/wheat consumption was independent of urban/rural identity. Chi-square p-values were adjusted with the BH method.

### Allele Frequencies at Lactase Persistence-Associated Variants

Since we did not sequence the host DNA in this project, we used published genetic data to estimate allele frequencies at positions known to be associated with lactase persistence. For the Sinhalese, Sri Lankan Tamil, Adivasi, and Spitian individuals, we used previously published data from the same communities^33,120,121^ (Table S9). For the Gond, Koya, Kani, and Kolam, who have either not been genome-sequenced previously or do not have available genotype data at the relevant positions (not covered on the genotyping array), we pooled a few proxy populations from the same geographic region (South and Central India) with linguistic overlap (Dravidian languages) and, in the case of the Gond, included genetically similar groups based on their published genetic affinities (Table S9)^107,122^. We used unpublished data for the Pochury from a study that is being prepared for submission (also co-led by corresponding authors MR and NR) and also considered them a proxy for the Mizo for whom there is limited genome-wide data available. For the Sinhalese, Sri Lankan Tamil, Adivasi, Gond, Koya, Kani, and Kolam, we estimated the allele frequency at the –13910 C/T (rs4988235) variant, commonly associated with lactase persistence in Europeans and many South Asians^87^. Since the Spitian and Pochury are genetically closely related to Tibetans/East Asians, we estimated allele frequencies at variants associated with populations from these regions^88^ in addition to –13910: –13838 G/A (rs1575359915), –13906 T/A (rs1679771596), and –13908 C/T (rs4988236).

## Author information

M.R. and N.R. conceived and supervised the project.

S.L.R. and N.P. designed and coordinated the experiments, with supervision from N.R. and M.R.

N.P., B.A., G.A., T.P.I., T.J., R.K., A.K., J.M.K., L., J.L., T.N., T.P., K.P., R.Ran., S.P.S., T.S., S.S., E.S., V.S., D.T., K.H.T., A.H.J.W., R.R.Y., and N.R. conducted participant recruitment and field sampling.

N.P., S.L.R., and V.B. performed data generation, with support from B.A., A.D., S.G., and R. and supervision from A.S., N.R., and M.R.

S.L.R. performed the principal components analyses, differential abundance analyses, co-abundance networks, taxon-phenotype associations, and taxon-diet associations, with supervision from M.R. R.Ram and R.J.A. performed the bioinformatic processing of sequencing reads and taxonomic classification, with supervision from A.S. and R.B. J.W.S. performed the SCALES analysis, with supervision from A.S.R. J.A.U.A. calculated the lactase persistence allele frequencies, with supervision from M.R.

S.L.R. and M.R. wrote the original draft of the manuscript, with input from R.J.A., V.B., R.B., E.R.D., T.P.I., N.P., J.W.S., and N.R. T.P.I., T.J., J.M.K., L., J.L., K.P., N.P., R.Ran, T.S., E.S., A.H.J.W., and R.R.Y. contributed community-specific ethnographic summaries.

All authors reviewed and approved the manuscript.

## Declaration of interests

The authors declare no competing interests.

## Data availability

Raw 16S reads are available on the Sequence Read Archive (SRA) under BioProject XYZ.

## Acknowledgements

We are deeply appreciative of the communities, community members, and donors whose openness, generosity, and willingness to participate made this research possible, and Adivasi leader Uruwarige Wannila Aththo and the clan leaders of Rathugala, Vakarei and Pollebedda, Sri Lanka. We acknowledge the Pochury Hoho, the various village councils, Chuthitu, the Chairman of the New Phor Village Council, and the Kohima Pochury people. We further acknowledge all individuals who directly or indirectly assisted in the field sample collection, especially Dr. Aswany Thomas (Medomas Healthcare, Thiruvananthapuram, India), Dr. M. Navas (KSCSTE-JNTBGRI, Thiruvananthapuram, India), Somaiah, Tekam Rukum Bhai, Tekam Ramu, Athram Mukund Rao, Madavi Bapu Rao, Tekam Jangu, Tanzin Angmo, Aman Sharma, Dr. Kishor Taram, Dr. Vijay Lekhami, Sunil Chichghare, Ramesh Sant, Sushil Kohad, the members of the Tualte Welfare Committee, and the Young Mizo Associations and Village Council members. We thank Thilina Pallethenna, Former Project Manager, Central Cultural Fund, Sri Lanka, for assisting in the collection of samples. We thank the University of Chicago Center for Research Informatics (USA) for providing access to high-performance computing resources. This project was funded by NIH Grant R35GM143094, Duchossois Family Institute Pilot Funding, University of Chicago, and University of Chicago start-up funds to M.R.; Birbal Sahni Institute of Paleosciences (BSIP) in-house project no. 7.3 to N.R; National Research Council, Sri Lanka, grant no. 17–042 to R.R.; NIH Grant R35GM146980 to E.R.D.; and NIH 5T32GM139782 to S.L.R. We are grateful to David Witonsky for helpful discussion and analytical support, Austin Rieger for technical support, and the Director of BSIP for supporting laboratory work in India.

## Declaration of generative AI and AI-assisted technologies in the manuscript preparation process

During the preparation of this work, the authors used ChatGPT (OpenAI) to assist with debugging analysis code and minor grammar editing.

## Notes

### Competing Interest Statement

The authors have declared no competing interest.

