## Supplementary Figures for "Distinct trajectories of urbanization shape the human gut microbiome across South Asia"

#### Gut microbial markers of lifestyle transition in 575 diverse South Asians

**This PDF file includes:**

Supplementary Text
Figs. S1 to S16
Tables S2 to S5, S7 to S9

**Other Supplementary Materials for this manuscript include the following:**

Data S1
Table S1, S6

**Table S1: Metadata and Diet Survey** **3**
**Table S2: Sampling Breakdown** **4**
**Table S3: Comparison of Shannon diversity in SAMBAR vs regional references** **5**
**Table S4: PERMANOVA test for statistical significance of regional clusters on the global PCA** **6**
**Table S5: ANOVA for beta dispersion of regional clusters on the global PCA** **7**
**Table S6: MaAsLin2 Results of differential abundance of taxa** **8**
**Table S7: Community distances from mean composition** **9**
**Table S8: Relative abundance of "main" CAGs vs smaller CAGs** **10**
**Table S9: Lactase persistence allele frequencies** **11**
**Figure S1: Principal Components Analysis of SAMBAR + 4 global cohorts** **12**
**Figure S2: Shannon Diversity by Region (SAMBAR vs Global Panel)** **13**
**Figure S3: All tested taxa, by world region** **14**
**Figure S4: Full Co-abundance network with SAMBAR-specific connections highlighted** **15**
**Figure S5: Full Co-abundance network with non-SAMBAR-exclusive connections highlighted** **16**
**Figure S6: Differential abundance heatmaps with non-Kolam communities as reference** **17**
Figure S6A: Gond 17
Figure S6B: Spitian 18
Figure S6C: Kani 19
Figure S6D: Koya 20
Figure S6E: Mizo 21
Figure S6F: Pochury 22
Figure S6G: SL Adivasi 23
Figure S6H: SL Tamil/Sinhalese 24
Figure S6I: All SAMBAR communities vs Sub-Saharan Africa 25
**Figure S7: Lifestyle PCA normalized by community** **26**
Figure7A: Normalized Lifestyle PCA colored by Urban/Rural 26
Figure7B: Normalized Lifestyle PCA colored by Community 26
**Figure S8: SCALES** **27**
**Figure S9: Megamonas abundance versus phenotypes and diversity** **28**
**Figure S10: Phenotypes and diversity by Urban / Rural** **29**
**Figure S11: Oscillospiraceae UCG-005 abundance vs phenotypes and diversity** **30**
**Figure S12: Full CAG plot for SAMBAR** **31**
**Figure S13: CAG relative abundance, merged by community-lifestyle** **31**
**Figure S14: CAG 6 vs Shannon Diversity** **32**
**Figure S15: CAG 5 vs Shannon Diversity** **33**
**Figure S16: Wheat/Yogurt in Urban/Rural** **34**
**Data File S1: Human Microbiome Compendium Subset** **35**

#### Table S1: Metadata and Diet Survey

(Separate csv file): Table of metadata for the 575 SAMBAR individuals analyzed in the study. Includes four categories of variables: demographic information (columns C:AA), phenotypes (columns AB:AJ), dietary survey (columns AK:BU), and sequencing metadata (columns BV:CE).

Table S2: Sampling Breakdown

| Table S1. Cohort demographics by community and lifestyle | | | | |
|---|---|---|---|---|
| | | N | Gender (M/F) | Age (Mean ± SD) |
| Gond | Rural | 53 | M=27, F=26 | 34 ± 9.3 |
| | Urban | 44 | M=20, F=24 | 40 ± 8.9 |
| Kani | Rural | 25 | M=6, F=19 | 42 ± 8.1 |
| | Urban | 42 | M=12, F=30 | 43 ± 7.2 |
| Kolam | Rural | 27 | M=15, F=12 | 36 ± 9.4 |
| | Urban | 38 | M=26, F=12 | 34 ± 11.3 |
| Koya | Rural | 30 | M=11, F=19 | 38 ± 7.1 |
| | Urban | 25 | M=7, F=18 | 40 ± 9.3 |
| Mizo | Rural | 30 | M=13, F=16 | 42 ± 11.6 |
| | Urban | 47 | M=22, F=22 | 44 ± 10.4 |
| Pochury | Rural | 29 | M=15, F=14 | 38 ± 6.6 |
| | Urban | 41 | M=27, F=14 | 35 ± 9.9 |
| Spitian | Rural | 25 | M=14, F=11 | 42 ± 10.6 |
| | Urban | 32 | M=10, F=22 | 36 ± 11.3 |
| Sri Lankan Adivasi | Rural | 42 | M=20, F=22 | 38 ± 9.6 |
| Sri Lankan Tamil / Sinhalese | Urban | 45 | M=26, F=19 | 38 ± 10.1 |

#### Table S3: Comparison of Shannon diversity in SAMBAR vs regional references

Differences in mean Shannon diversity between SAMBAR and each global reference cohort.

| comparison | mean_SAMBAR | mean_comparison | mean_diff | t_statistic | p_value |
|---|---|---|---|---|---|
| **Eastern and South-Eastern Asia** | 3.0054 | 2.4509 | -0.5546 | 18.9269 | 1.64E-68 |
| **Europe and Northern America** | 3.0054 | 2.5104 | -0.495 | 13.813 | 2.62E-39 |
| **Northern Africa and Western Asia** | 3.0054 | 2.6057 | -0.3997 | 13.8792 | 3.17E-40 |
| **Sub-Saharan Africa** | 3.0054 | 2.8259 | -0.1795 | 5.4984 | 4.97E-08 |

Table S4: PERMANOVA test for statistical significance of regional clusters on the global PCA

PERMANOVA on the coordinates for SAMBAR and the four regions on PC 2 and 3.

| | Df | SumOfSqs | R2 | F | Pr(>F ) |
|---|---|---|---|---|---|
| **Model** | 4 | 1550.178 | 0.334 | 360.092 | 0.001 |
| **Residual** | 2870 | 3088.799 | 0.666 | NA | NA |
| **Total** | 2874 | 4638.977 | 1 | NA | NA |

Table S5: ANOVA for beta dispersion of regional clusters on the global PCA

Significant differences in the dispersion or 'tightness' of the five cohorts on the global PCA

| | Df | Sum Sq | Mean Sq | F value | Pr(>F) |
|---|---|---|---|---|---|
| **Groups** | 4 | 16.427 | 4.107 | 19.69 | 5.283E-16 |
| **Residuals** | 2870 | 598.618 | 0.209 | NA | NA |

Table S6: MaAsLin2 Results of differential abundance of taxa

(Separate Excel file); outputs from MaAsLiN2 (“results.tsv”) as described in main text.

Table S7: Community distances from mean composition

We calculated the mean microbial composition of the full SAMBAR dataset. We then calculated the mean composition for each community separately. Both mean compositions were centered-log-ratio-transformed and the Euclidean distance between each community mean and the SAMBAR mean was calculated.

| | sort(dist_to_mean) |
|---|---|
| **Kolam** | 5.0156 |
| **Gond** | 5.69601 |
| **Koya** | 7.9408 |
| **SL_Adivasi** | 11.2492 |
| **Mizo** | 11.9735 |
| **Pochury** | 12.3448 |
| **Spitian** | 12.6833 |
| **SLTamil_Sinhalese** | 13.0484 |
| **Kani** | 13.3519 |

Table S8: Relative abundance of “main” CAGs vs smaller CAGs

All individuals were summed at the community level, and abundances of CAGs were summed: CAGs 1:6 (the “primary” CAGs discussed; CAGs 7:10 (significantly less abundant CAGs), and OTHER (all taxa that were not assigned to a CAG)

| Community_Lifestyle | CAGs_1_to_6 | CAGs_7_to_10 | OTHER |
|---|---|---|---|
| Gond-Rural | 0.880 | 0.032 | 0.089 |
| Gond-Urban | 0.892 | 0.033 | 0.074 |
| Spitian-Rural | 0.872 | 0.027 | 0.101 |
| Spitian-Urban | 0.903 | 0.017 | 0.080 |
| Kani-Rural | 0.896 | 0.032 | 0.073 |
| Kani-Urban | 0.888 | 0.039 | 0.073 |
| Kolam-Rural | 0.860 | 0.024 | 0.116 |
| Kolam-Urban | 0.854 | 0.052 | 0.093 |
| Koya-Rural | 0.889 | 0.041 | 0.069 |
| Koya-Urban | 0.885 | 0.041 | 0.074 |
| Mizo-Rural | 0.835 | 0.089 | 0.076 |
| Mizo-Urban | 0.874 | 0.066 | 0.060 |
| Pochury-Rural | 0.896 | 0.040 | 0.064 |
| Pochury-Urban | 0.878 | 0.061 | 0.061 |
| SriLankan-Rural | 0.872 | 0.025 | 0.103 |

#### Table S9: Lactase persistence allele frequencies

We screened for the presence of known lactase persistence variants in members of the same community, when possible, or in anthropologically/geographically and genetically informed proxy populations with expected similarities in lactase persistence.

| **Community** | **Variant** | **Frequency** | **Community Notes** |
|---|---|---|---|
| Sinhalese | -13910 C/T (rs4988235) | 0/70 | Urban Aragon et al 2025 |
| Adivasi | -13910 C/T(rs4988235) | 1/38 | Urban Aragon et al 2025 |
| Sri Lankan Tamil | -13910 C/T (rs4988235) | 3/70 | 1000 Genomes |
| Gond, Koya, Kani, Kolam | -13910 C/T (rs4988235) | 0/74 | Proxy populations: GenomeAsia 100K: Abujmaria, Muria, BisonHornMaria, Paniya |
| Pochury | -13910 C/T (rs4988235) | 0/40 | Unpublished data |
| Pochury | -13838 G/A (rs1575359915) | 0/36 | Unpublished data |
| Pochury | -13906 T/A (rs1679771596) | 0/40 | Unpublished data |
| Pochury | -13908 C/T (rs4988236) | 0/40 | Unpublished data |
| Spitian | -13910 C/T (rs4988235) | 0/40 | Bandyopadhyay et al 2025 |
| Spitian | -13838 G/A (rs1575359915) | 1/40 | Bandyopadhyay et al 2025 |
| Spitian | -13906 T/A (rs1679771596) | 0/40 | Bandyopadhyay et al 2025 |
| Spitian | -13908 C/T (rs4988236) | 0/40 | Bandyopadhyay et al 2025 |

### Figure S1: Principal Components Analysis of SAMBAR + 4 global cohorts

S1A: Principal Components Analysis (Aitchison PCA) of SAMBAR cohort + 4 global regions (as described in Figure 2A). S1B: PC 1 and 2 of SAMBAR + 4 regions, colored by Shannon diversity (calculated at genus level). S1C: PC 2 and 3 of SAMBAR + 4 regions, with taxa contributing to variance

S1A

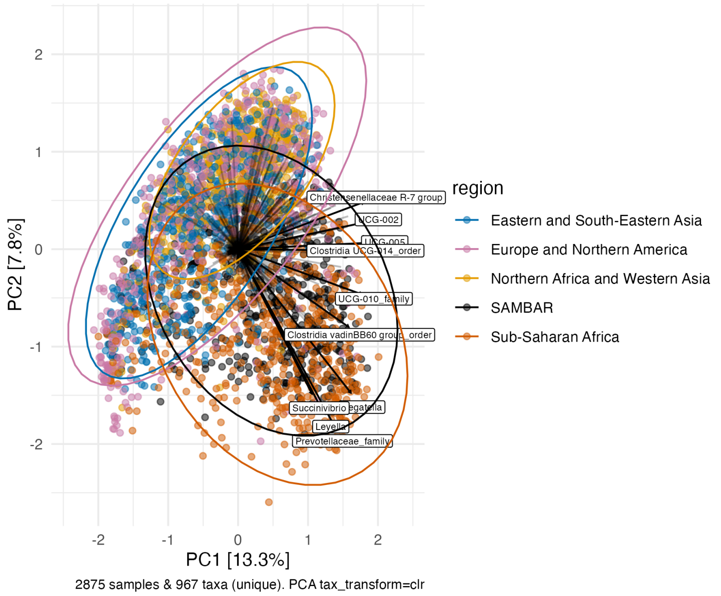


S1B

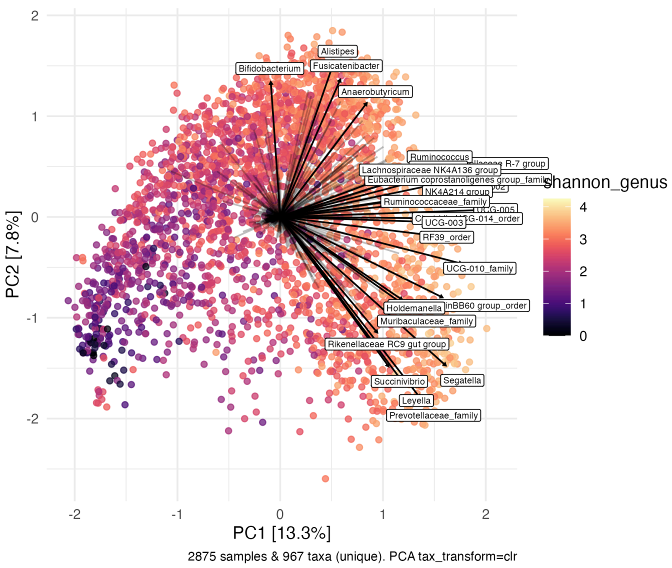


S1C

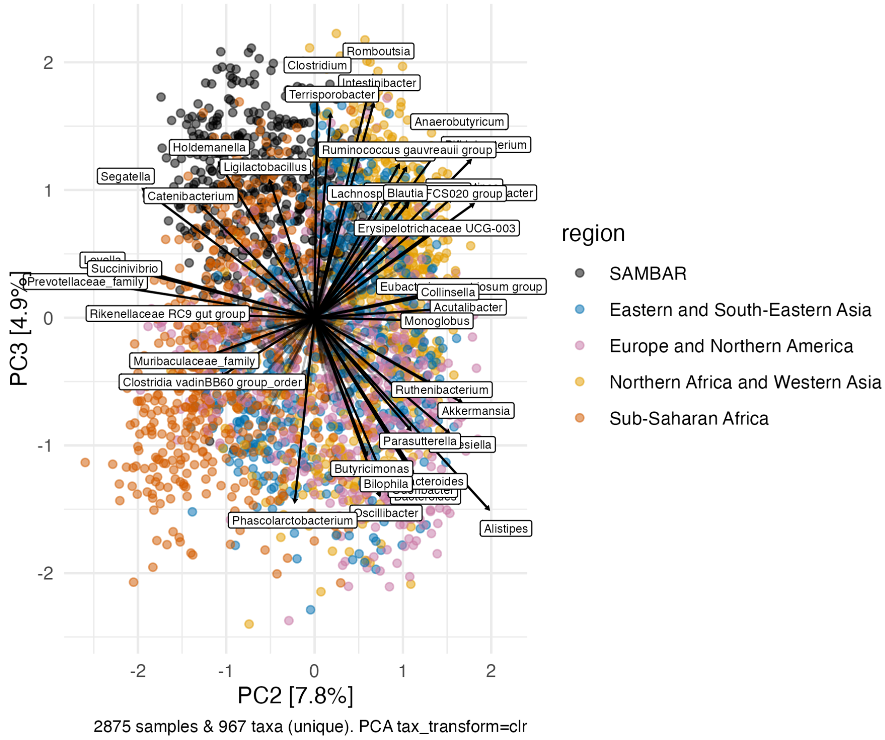

#### Figure S2: Shannon Diversity by Region (SAMBAR vs Global Panel)

Shannon diversity by region; results of significance testing when comparing SAMBAR with all global references are in Table S3.

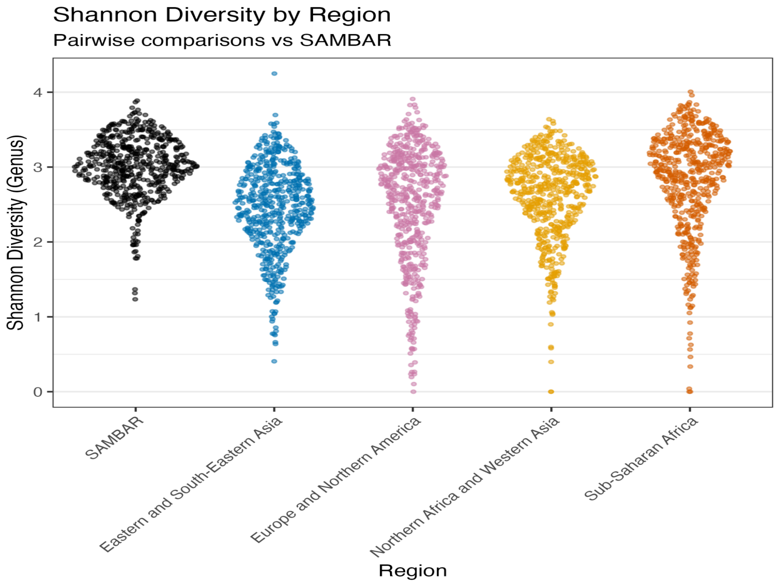

### Figure S3: All tested taxa, by world region

Extended version of Figure 2C. Green taxa are more abundant in SAMBAR; purple taxa are more abundant in the comparison cohort than in SAMBAR. Taxa are ordered from highest to lowest relative abundance in SAMBAR, represented by the black bars.

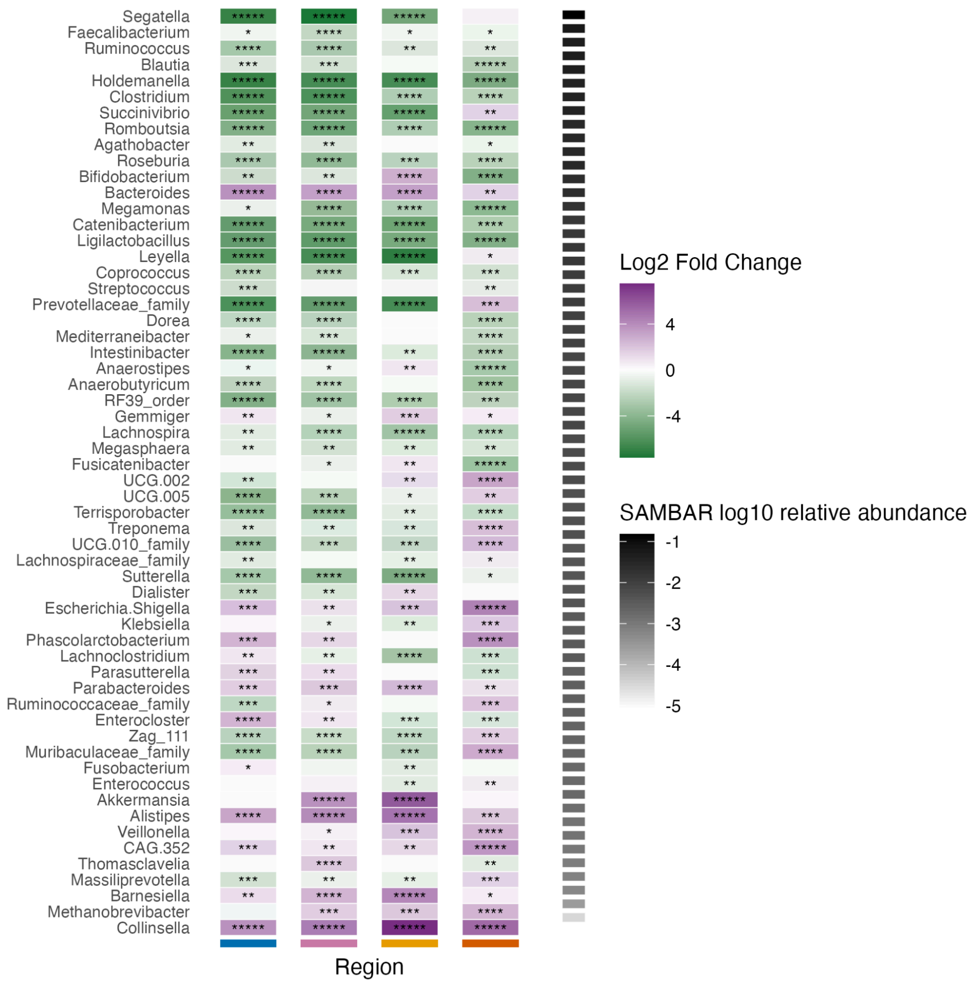

#### Figure S4: Full Co-abundance network with SAMBAR-specific connections highlighted

Extended version of network diagram in Figure 2D.

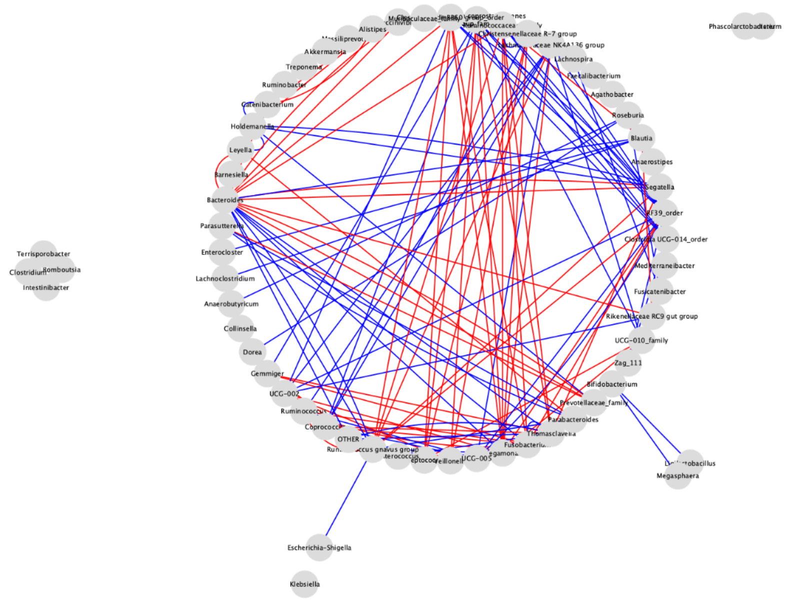

Figure S5: Full Co-abundance network with non-SAMBAR-exclusive connections highlighted

Extended version of network diagram in Figure 2E.

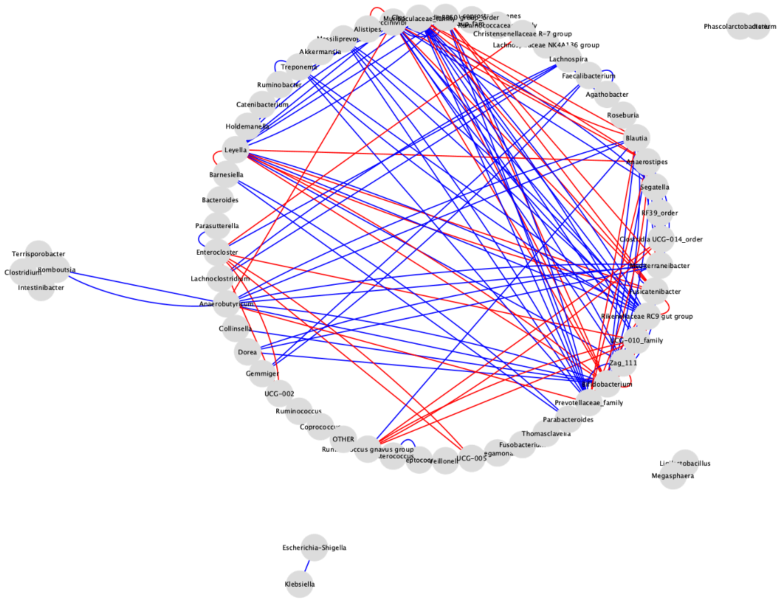

#### Figure S6: Differential abundance heatmaps with non-Kolam communities as reference

Differentially abundant taxa between each community within SAMBAR and all other communities in SAMBAR, controlling for lifestyle. Green taxa are more abundant in the comparison community; purple taxa are more abundant in the target community. Taxa are ordered by relative abundance in the target community, represented by black bars.

##### Figure S6A: Gond

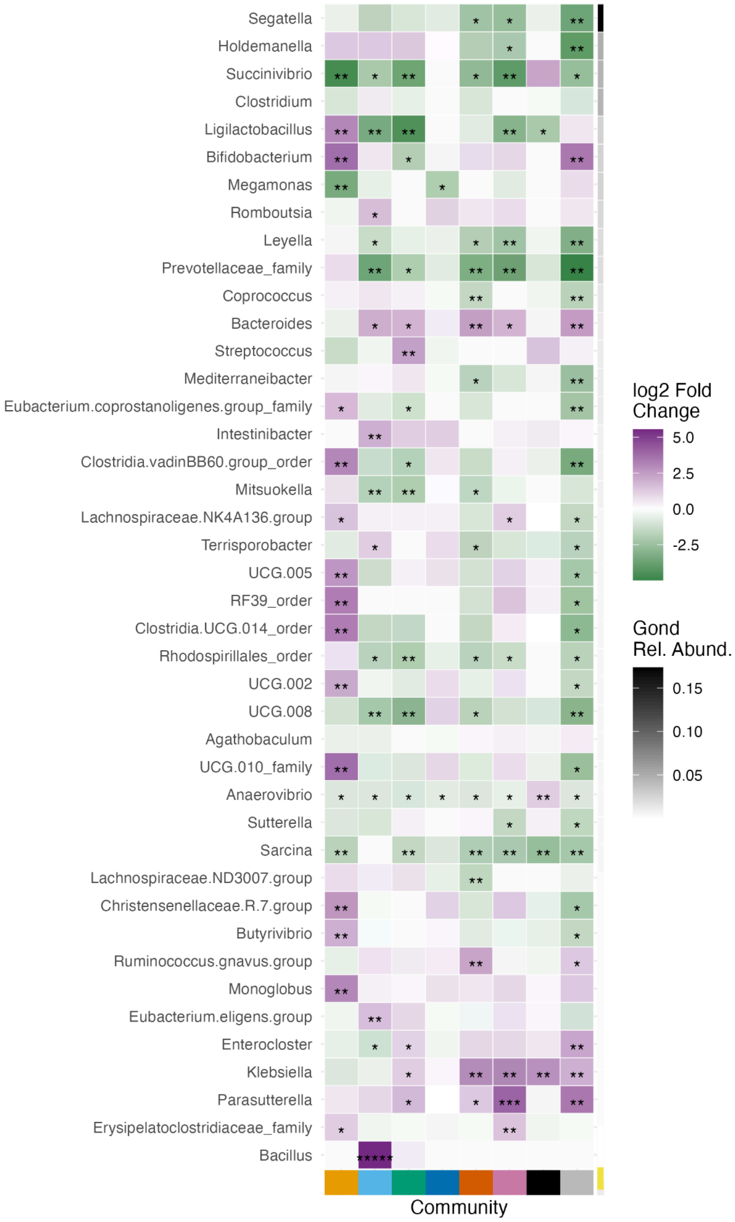

Figure S6B: Spitian

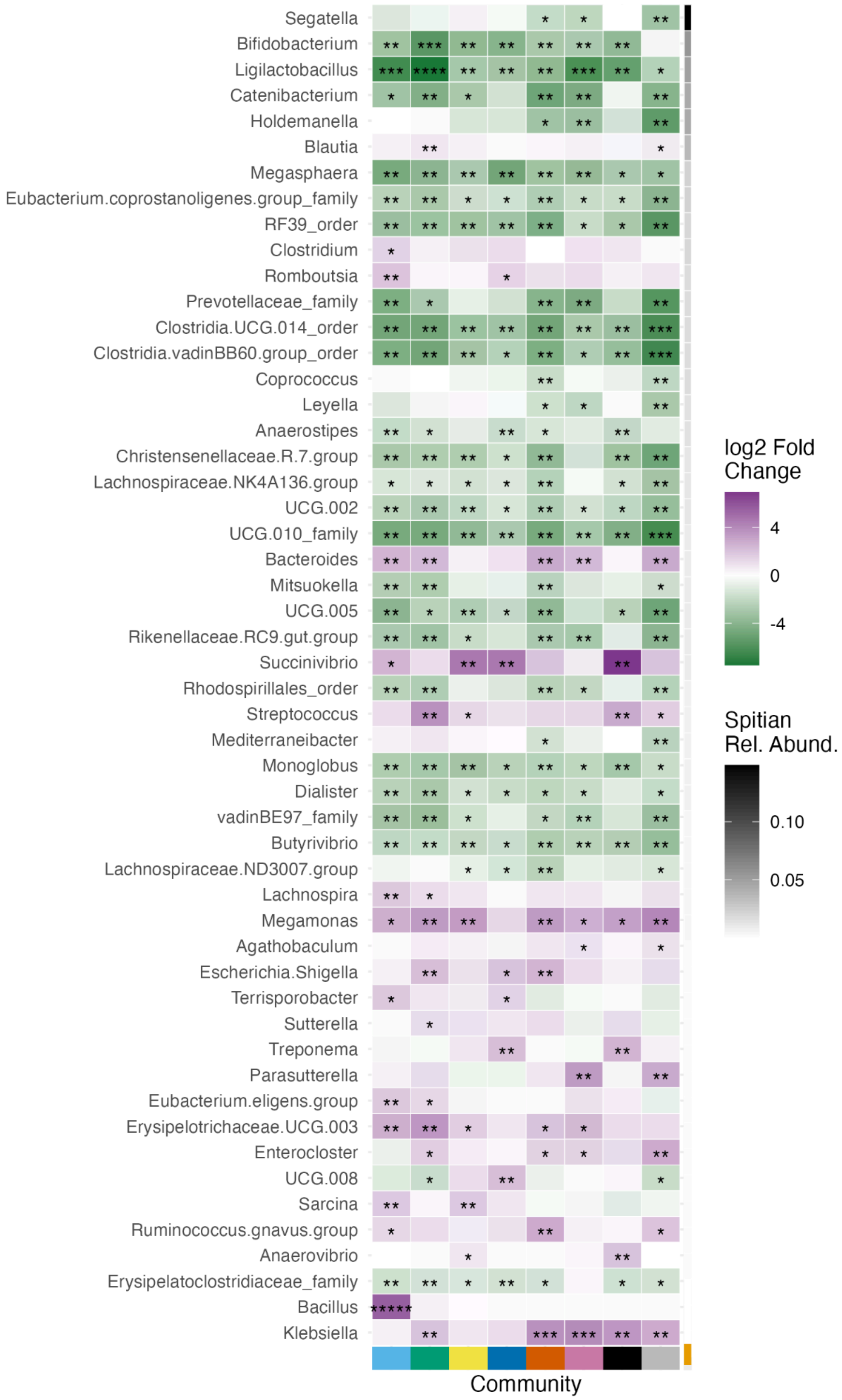

Figure S6C: Kani

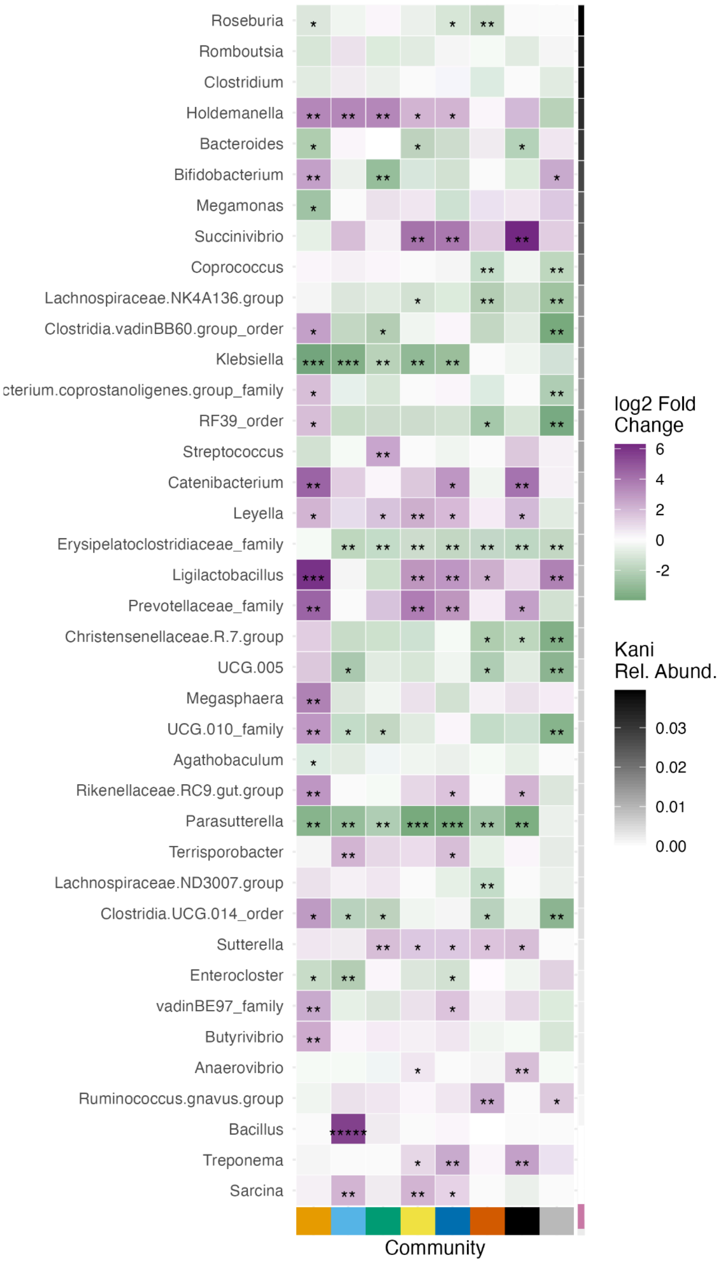

Figure S6D: Koya

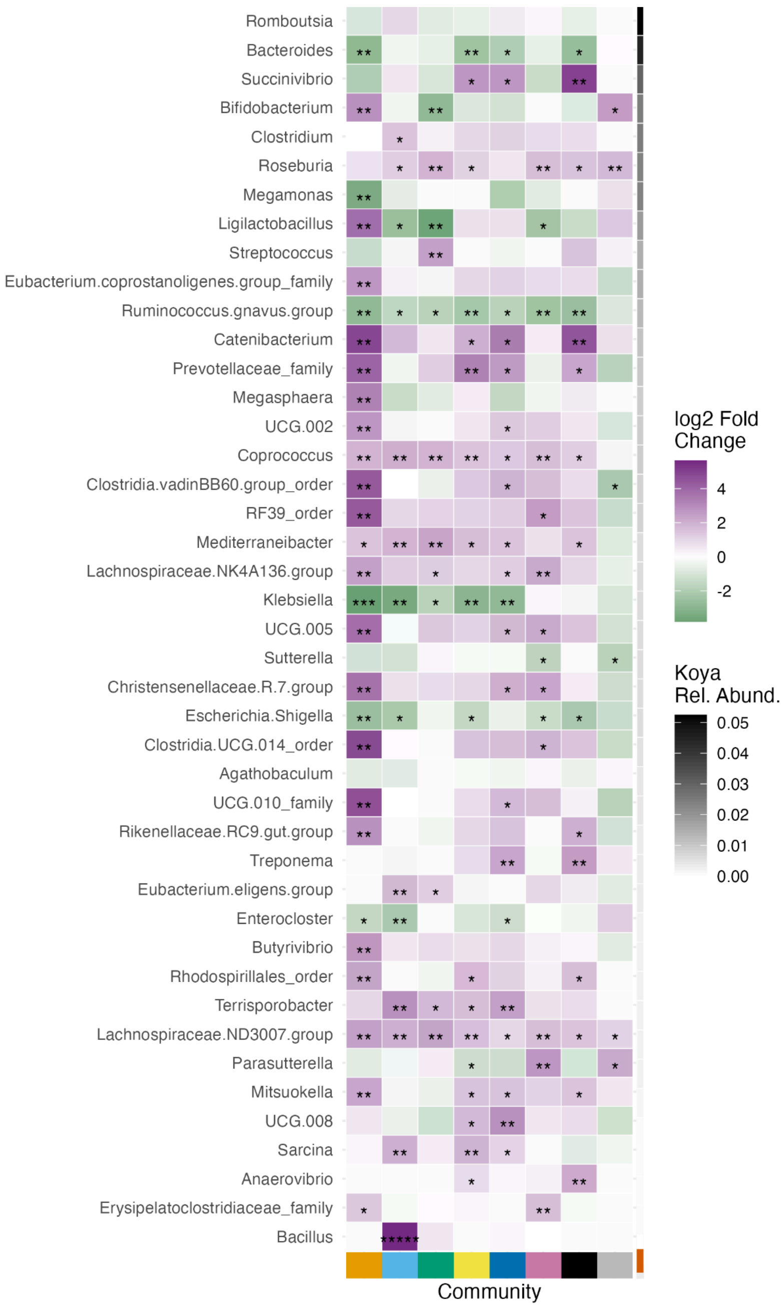

Figure S6E: Mizo

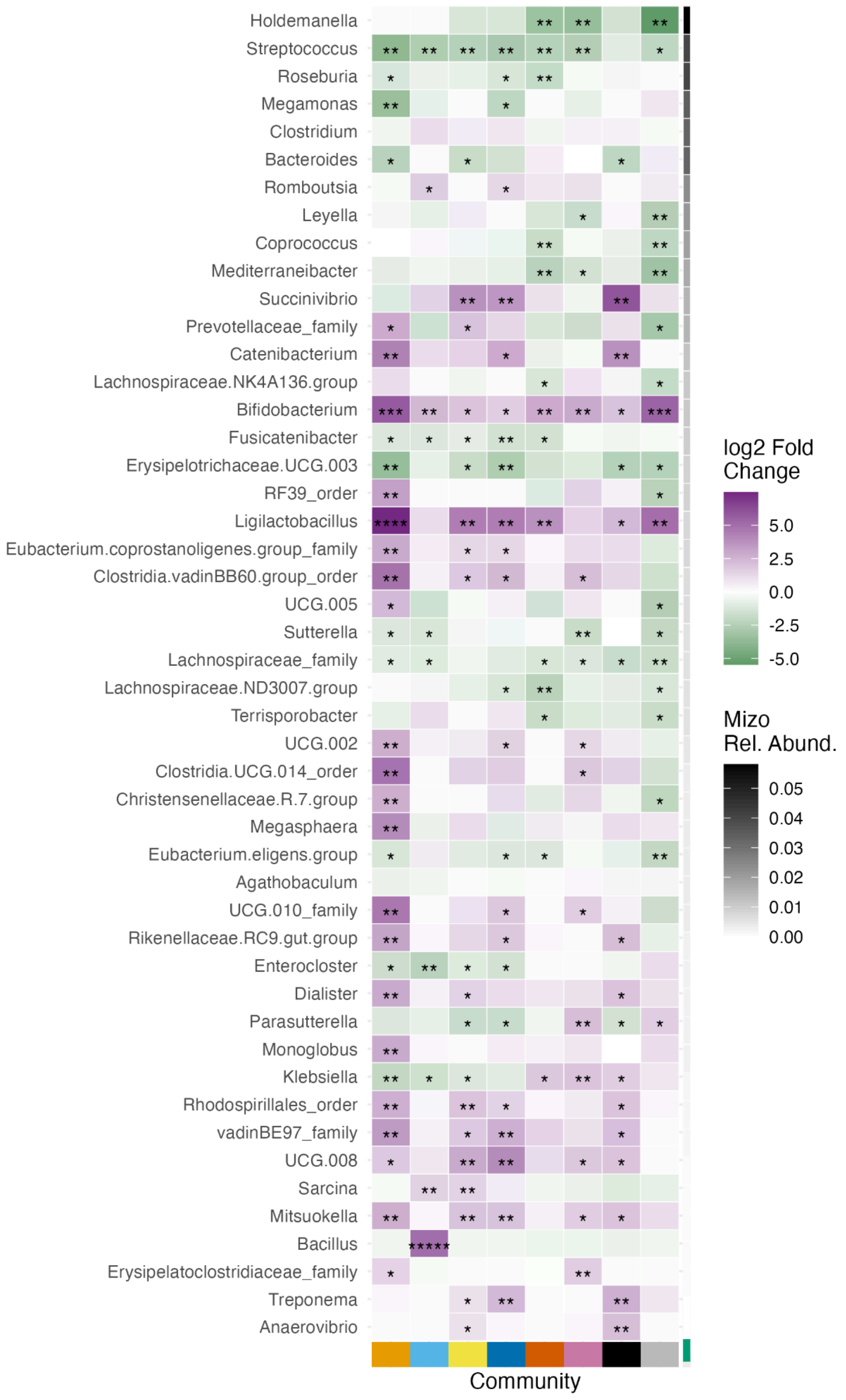

Figure S6F: Pochury

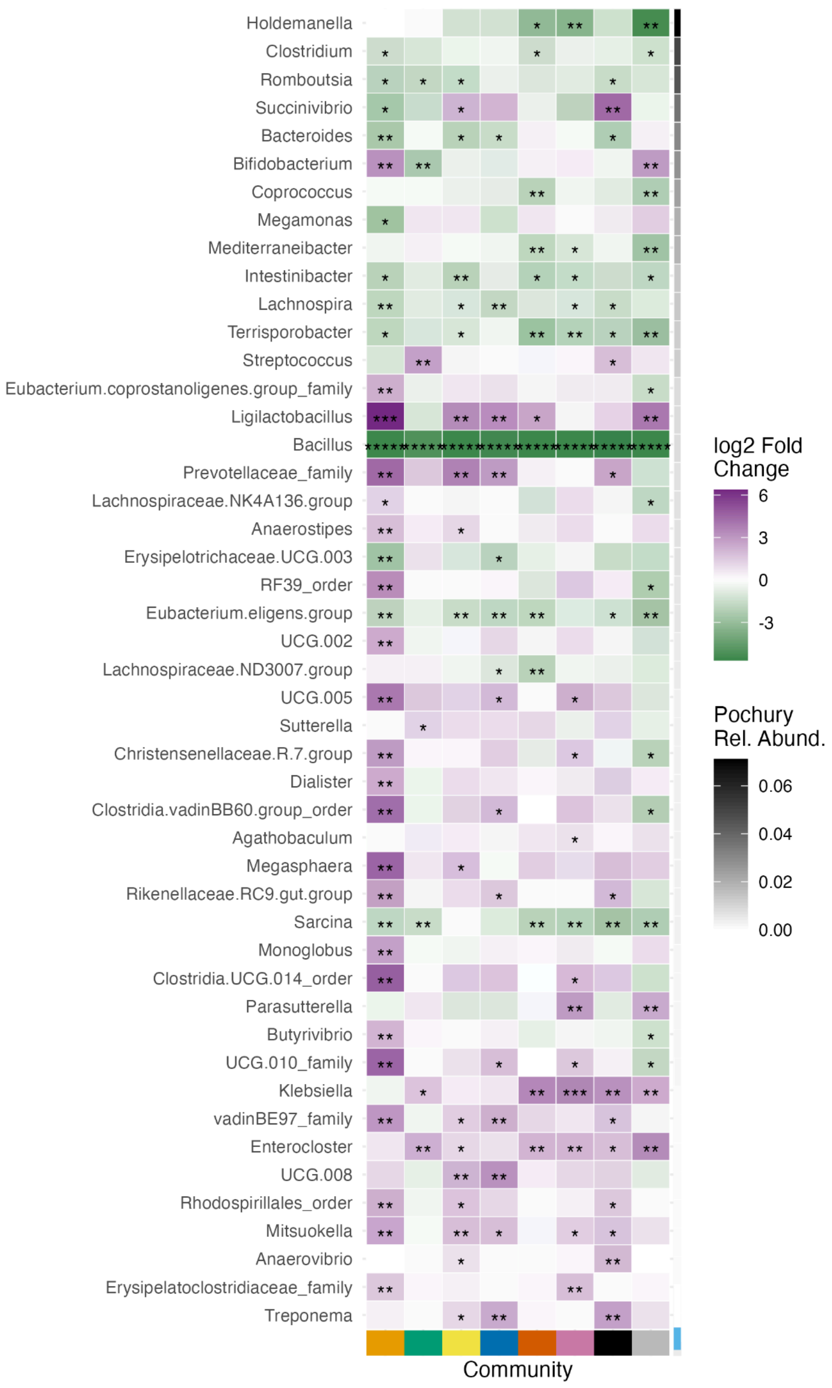

Figure S6G: SL Adivasi

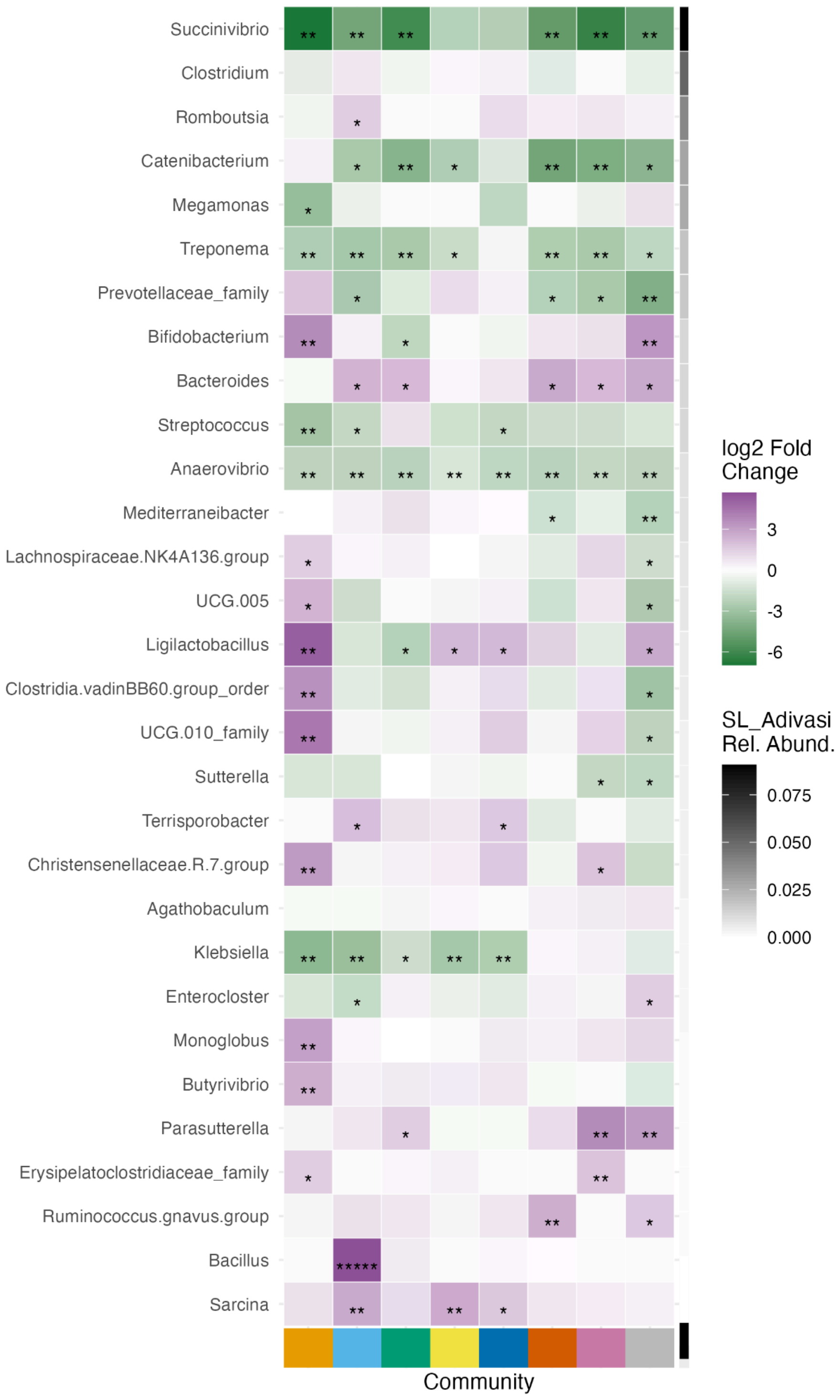

Figure S6H: SL Tamil/Sinhalese

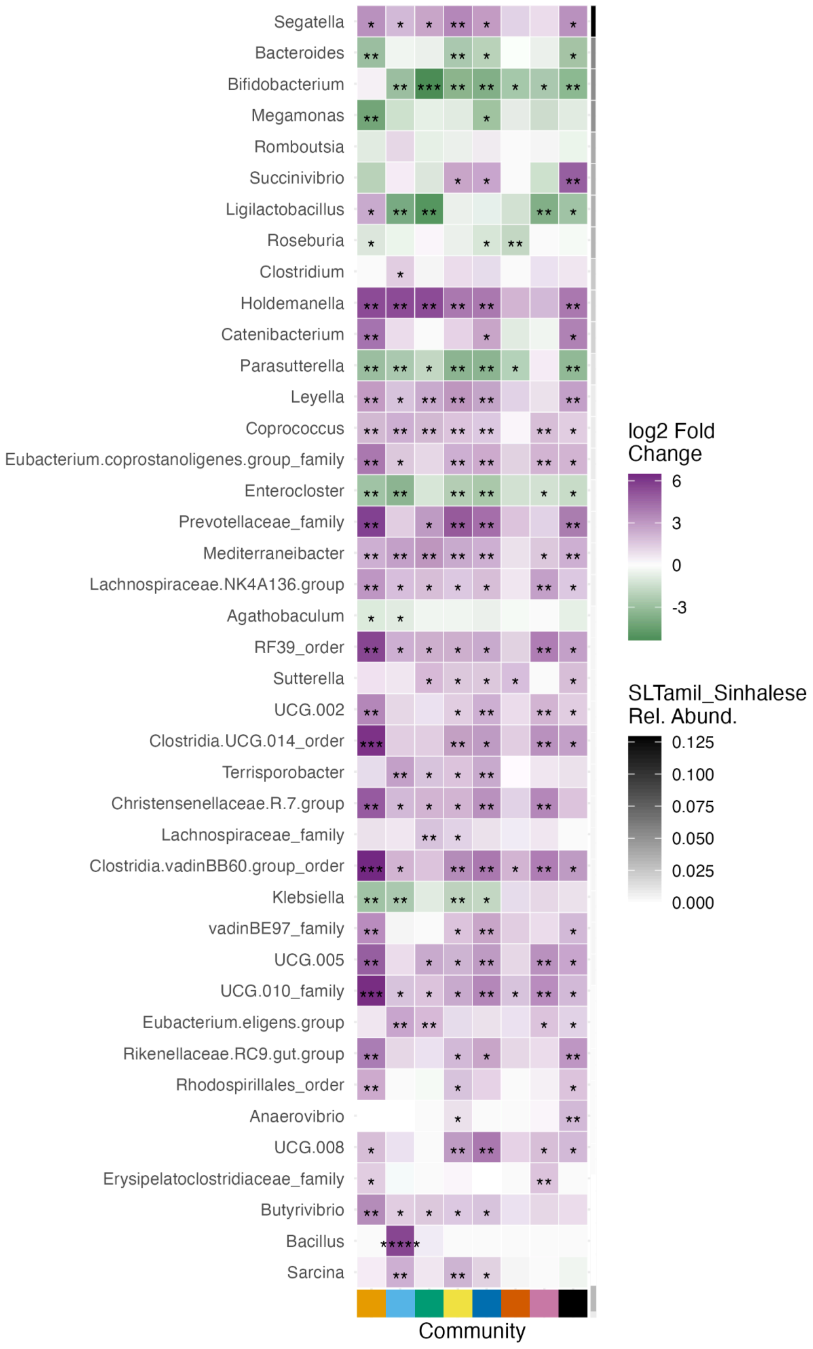

Figure S6I: All SAMBAR communities vs Sub-Saharan Africa

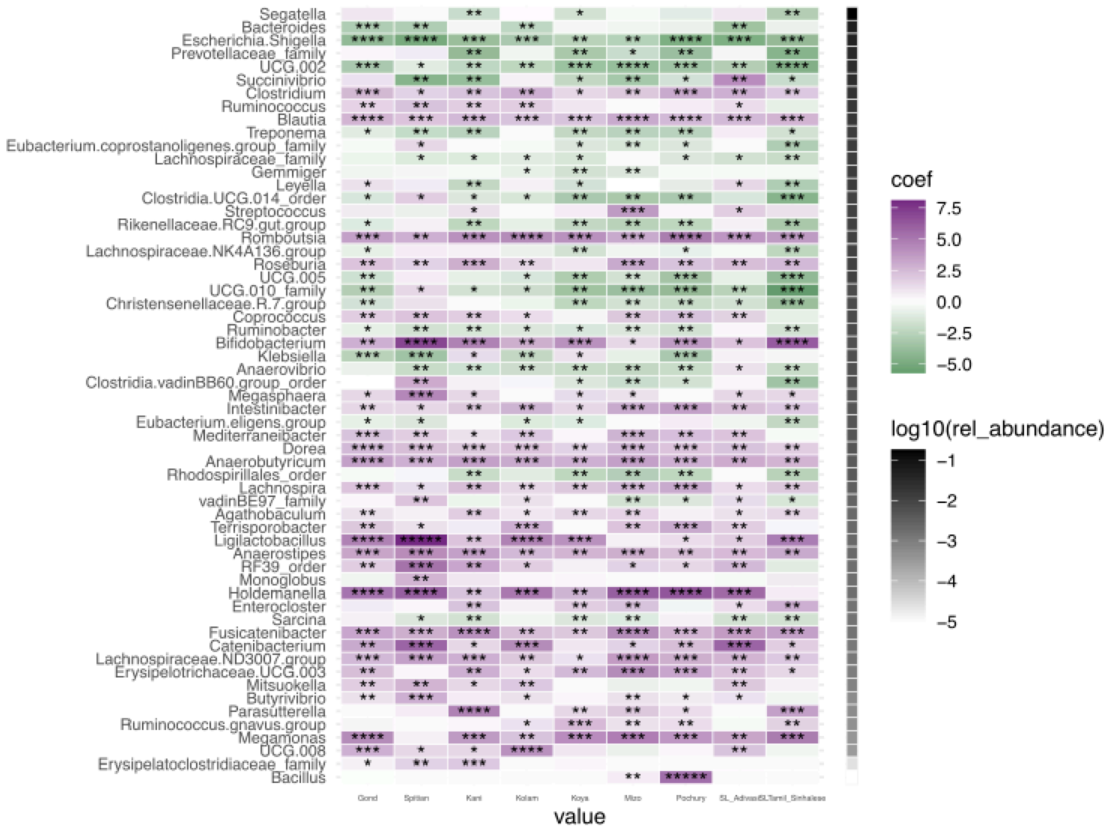

#### Figure S7: Lifestyle PCA normalized by community

Re-plotting of the PCA in Fig 4E. The mean coordinates for each community were calculated, and the per-community mean coordinates were subtracted from the coordinates for each point in the community.

##### Figure7A: Normalized Lifestyle PCA colored by Urban/Rural

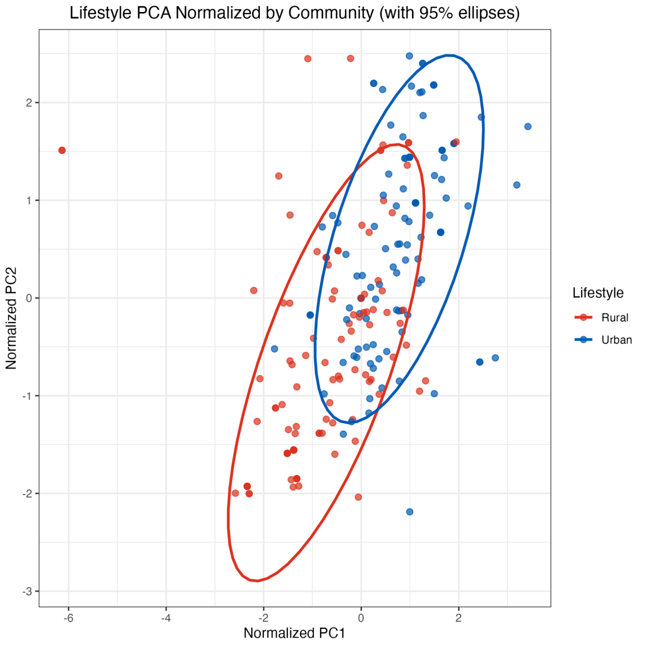


##### Figure7B: Normalized Lifestyle PCA colored by Community

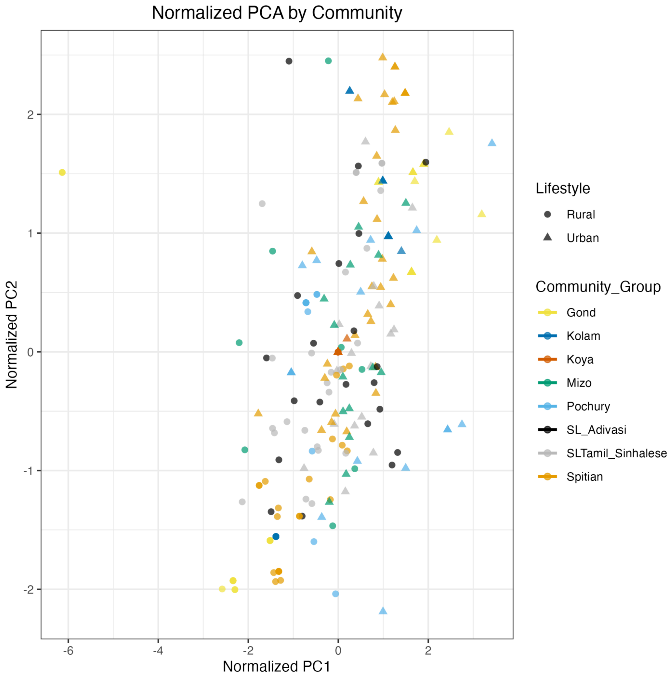

Figure S8: SCALES

Mutual Information (MI) box plots from SCALES

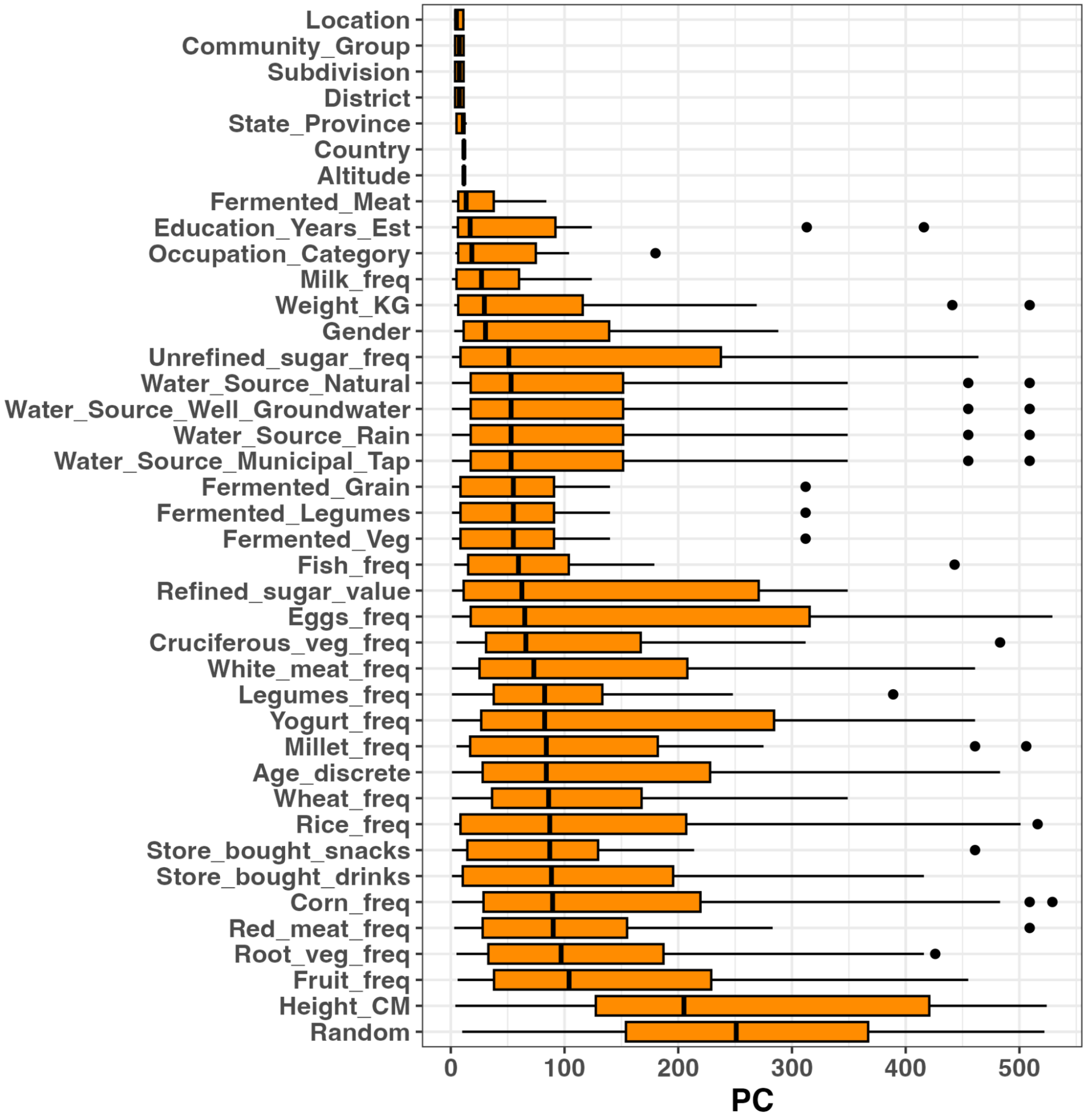

### Figure S9: Megamonas abundance versus phenotypes and diversity

Simple linear model of log1p(Megamonas abundance) vs quantitative phenotypes of interest and Shannon diversity in SAMBAR cohort

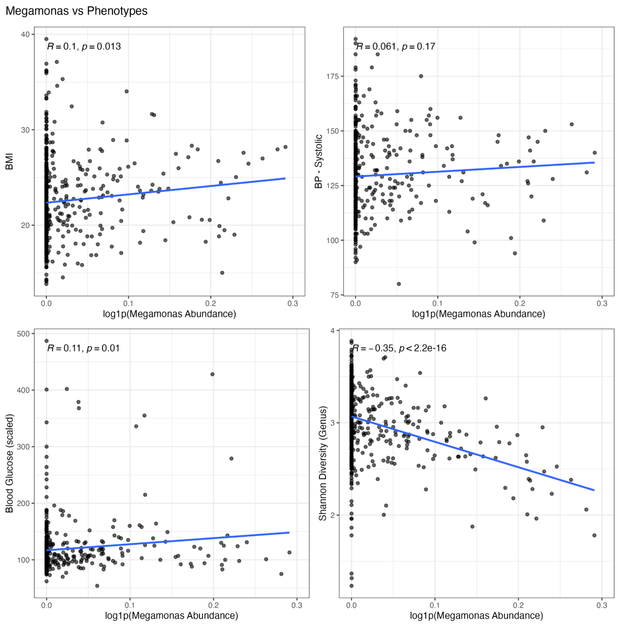

#### Figure S10: Phenotypes and diversity by Urban / Rural

Phenotype of interest and Shannon diversity plotted by Urban and Rural lifestyle - partial residuals calculated after controlling for Age, Gender, read count, and community group membership.

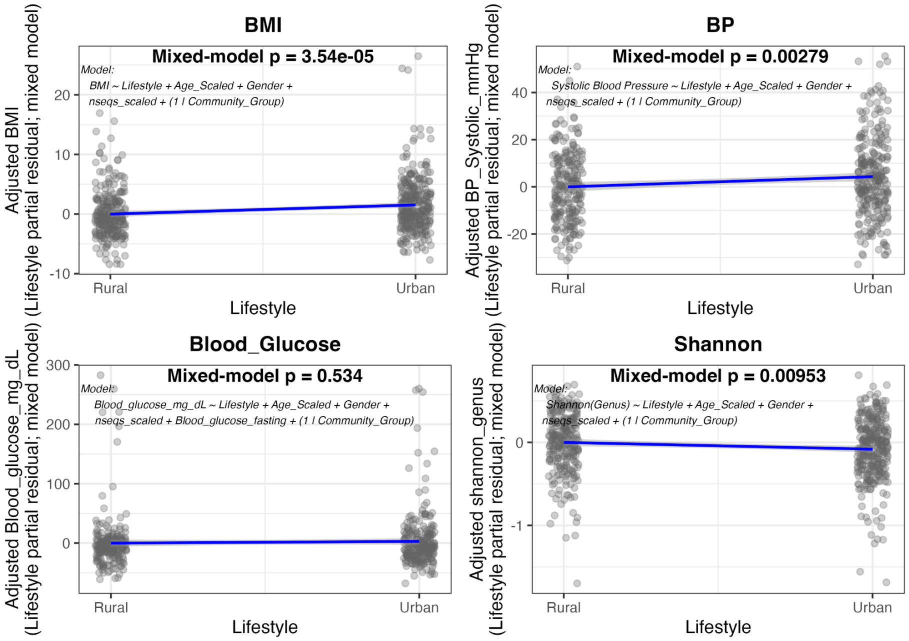

#### Figure S11: Oscillospiraceae UCG-005 abundance vs phenotypes and diversity

Simple linear model of log1p(UCG-005 abundance) vs quantitative phenotypes of interest and Shannon diversity in SAMBAR cohort

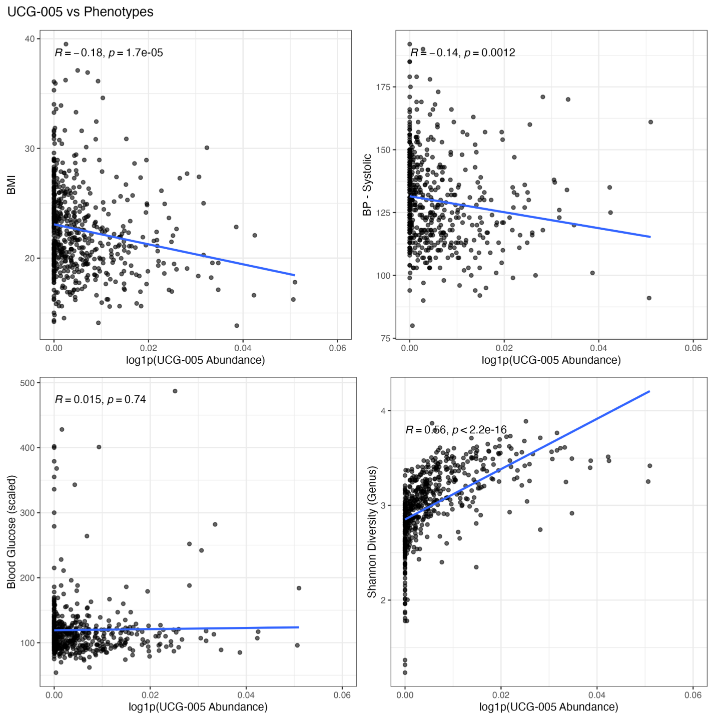

#### Figure S12: Full CAG plot for SAMBAR

Full plot for subset in Fig. 4D.

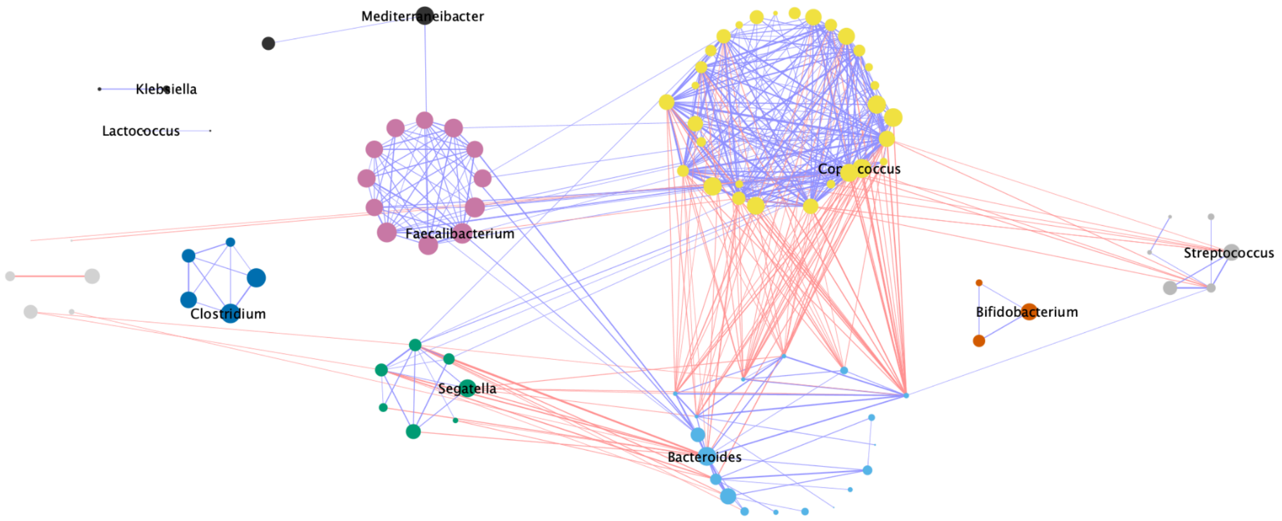


#### Figure S13: CAG relative abundance, merged by community-lifestyle

Samples were aggregated at the community-lifestyle level, and relative abundances for all taxa in each CAG in each aggregate-sample were plotted.

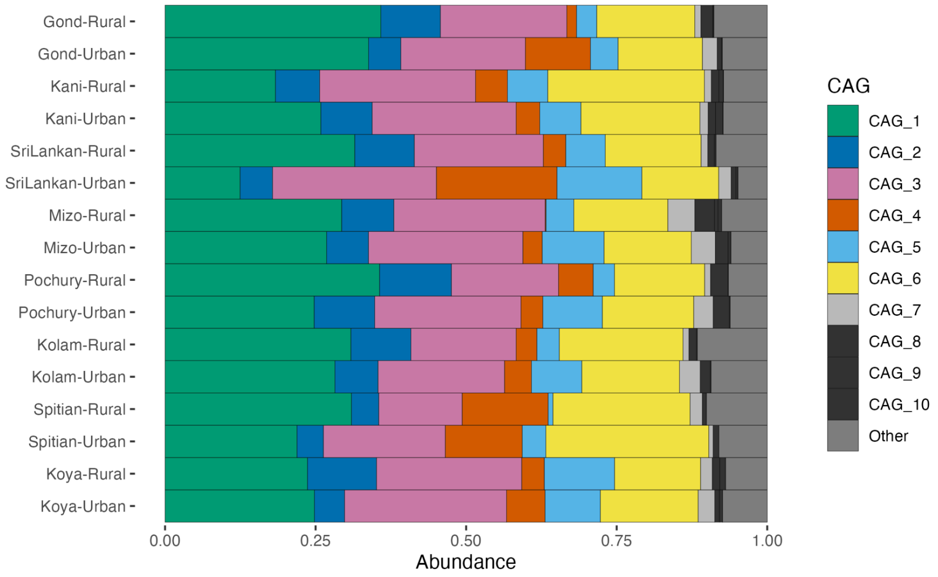

#### Figure S14: CAG 6 vs Shannon Diversity

Scatter plot of CAG 6 (UCG-005) abundance vs Shannon diversity (calculated at genus level) in SAMBAR cohort

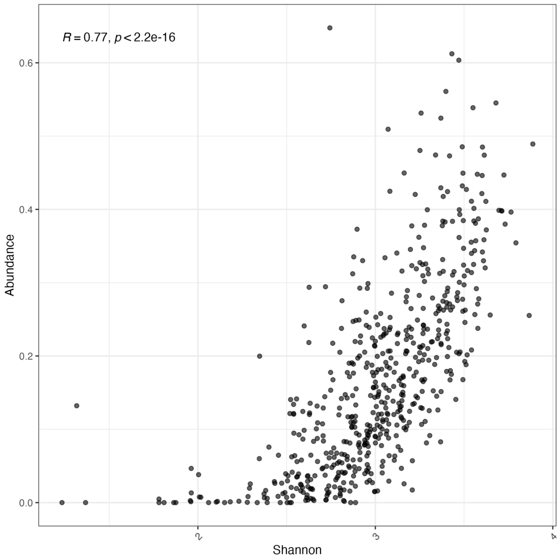

### Figure S15: CAG 5 vs Shannon Diversity

Scatter plot of CAG 5 (Bifidobacterium) abundance vs Shannon diversity (calculated at genus level) in SAMBAR cohort

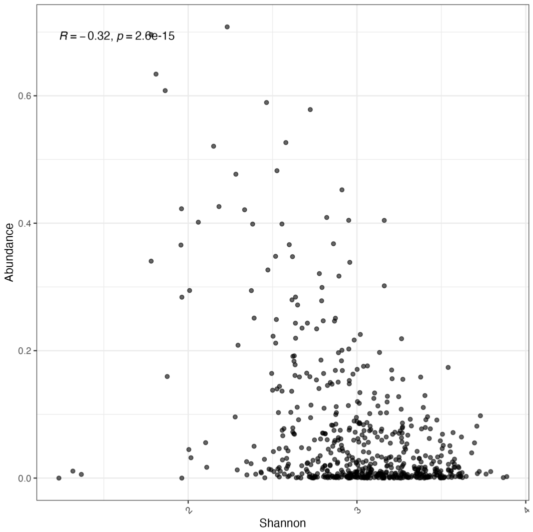

Figure S16: Wheat/Yogurt in Urban/Rural

Participants were categorized as either “Urban” or “Rural” and either “High” or “Low” consumers of wheat and yogurt. For each food product, a chi-square test for independence was performed between urban/rural status and high/low consumption. Significant results (in red) reflect communities where increased wheat or yogurt consumption is associated with urban lifestyle.

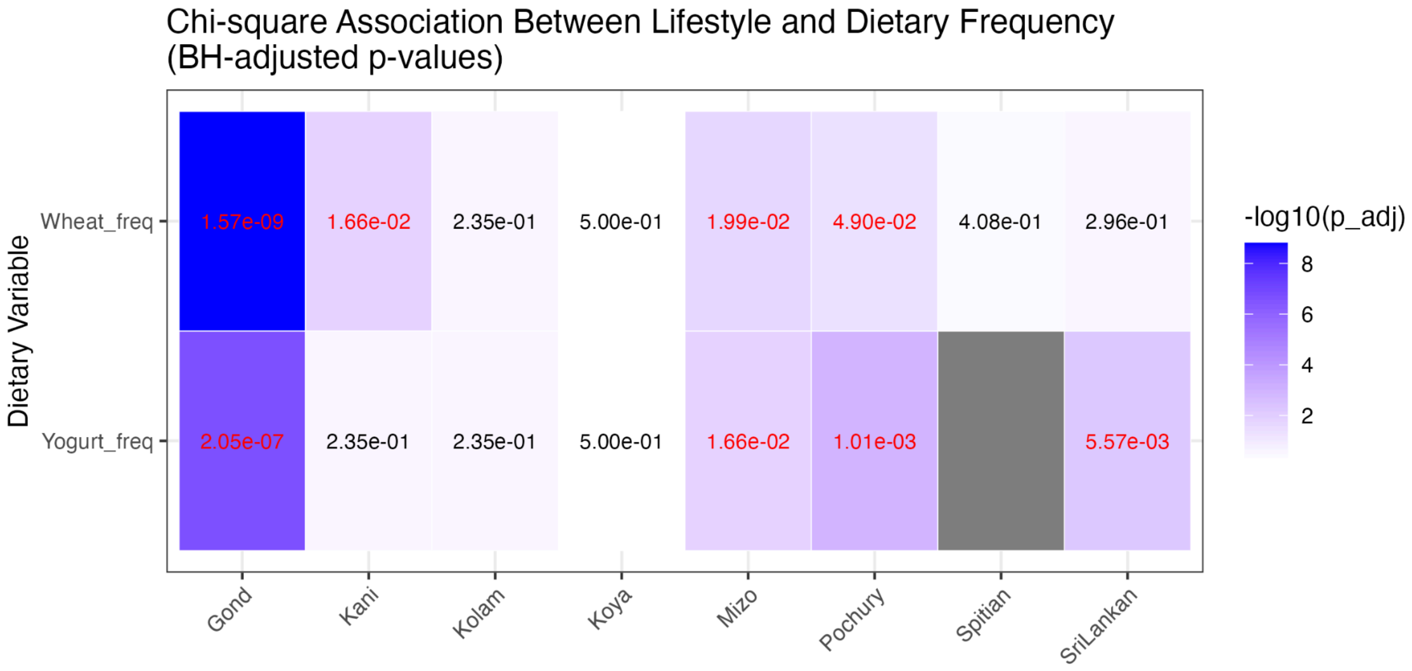

#### Data File S1: Human Microbiome Compendium Subset

(Separate text file); List of Human Microbiome Compendium individuals included in this study (575 per region, randomly selected from filtered compendium). File contains SRR ID for each individual.
